# Identifying and targeting abnormal mitochondrial localization associated with psychosis

**DOI:** 10.1101/2025.10.08.676630

**Authors:** Marzieh Haghighi, Donna McPhie, Mohammad Rohban, Erin Weisbart, David J. Logan, Kyle W. Karhohs, Suzann M. Babb, Jessica D. Ewald, Johan Fredin Haslum, Caitlin Ravichandran, Beth A. Cimini, Shantanu Singh, Bruce M. Cohen, Anne E. Carpenter

## Abstract

Therapeutics working by novel mechanisms are needed for patients with psychiatric conditions. Cell-based assays to identify candidates that reverse observed abnormalities, even if in subsets of patients, could accelerate the process. Here, we imaged peripheral cells (skin fibroblasts) of 168 individuals, stained for DNA, actin, and mitochondria. We found mitochondria tend to be farther from the cell border for patients who experience psychosis (including subsets of individuals with bipolar disorder, schizophrenia, and schizoaffective disorder), a phenotype not seen in patients diagnosed with major depression. Because the phenotype could be identified by a single metric, we could query existing databases of cells stained for their mitochondria and treated with various chemical or genetic perturbations. We identified compounds and genes both negatively and positively affecting the psychosis-associated phenotype, including some known to impact psychiatric conditions. Developing therapeutics with novel mechanisms is a complex multi-step challenge. This cell-based assay holds promise for virtual and physical screening to identify candidates for treating psychiatric conditions.

## Introduction

Many patients presenting with psychosis respond only partially or poorly to current treatments. They are in critical need of medicines with novel mechanisms of action, both to improve efficacy and reduce disabling side effects relative to existing medicines (Pereira et al. 2018; Stępnicki et al. 2018; Cohen and Öngür 2018; McIntyre et al. 2020; Kuperberg et al. 2021). These conditions include schizophrenia, schizoaffective disorder and bipolar disorder, which collectively have an enormous economic, social, and emotional impact (Bessonova et al. 2020; Kadakia et al. 2022; Lin et al. 2022).

Unlike many diseases, target-based drug discovery is not ideal for these conditions. Hundreds and even thousands of genes underlie the risk for psychotic disorders, and no single gene determines more than a modest proportion of risk (Trubetskoy et al. 2022). There is limited knowledge about the identity and function of all the proteins and other cellular components involved in the cellular processes that ultimately underlie and lead to psychosis. Current standard medications were discovered serendipitously. Predicting new protein or cellular targets whose chemical modulation will produce the desired antipsychotic effect in humans, based on existing understanding, is error-prone, making the unbiased approach of phenotypic assays appealing. However, ideal phenotypic assays involve a faithful, high-throughput model of the disease pathogenicity in a dish. This is inherently challenging for conditions involving the brain. Although complex brain organoids can be engineered (Qian et al. 2019; Tan et al. 2021), they are not well-suited to high throughput screening. More importantly, it is unclear what phenotypic effect ought to be sought in cells or organoids in order to identify compounds that relieve neuropsychiatric symptoms. Often, promising therapeutic hypotheses fail in the clinic given complexities at the organism level and a faulty link between a proposed target or phenotype and the human condition.

Rather than starting from a hypothesized phenotype based on existing scientific knowledge for a condition, recent evidence demonstrates that phenotypes can be identified in an unbiased way directly from patient samples, as we discuss next. Association of a cell phenotype with a condition does not prove causation; nevertheless, it enables drug screening, such that compounds reversing cell-based phenotypes are worth testing to see if they effectively reverse disease symptoms.

Patient fibroblasts, derived from a skin biopsy, are an appealing source material for identifying phenotypes because they are a cell type that is relatively accessible, acquirable from living patients, and sufficiently propagated for high-throughput screens. Although some fibroblasts share an origin with neuroectoderm, they would not be expected to have all of the necessary cellular pathways and structures to accurately reflect phenotypes for all human conditions. Still, phenotypes for several neurological conditions have been detected in patient fibroblasts (Olesen et al. 2022). Studies that revealed mitochondrial morphology phenotypes as compared to controls include fibroblasts from 41 patients with mid-stage idiopathic Parkinson’s disease (Antony et al. 2020), in several genetic and chemical neuronal models of Parkinson’s disease (Thomas et al. 2023), and in 46 patients with Parkinson’s – 32 sporadic, 14 carrying particular mutations (Schiff et al. 2022). A change in several mitochondrial functions was found in 100 sporadic Parkinson’s patients and 50 age-matched controls (Carling et al. 2020); mitochondrial count and mitochondrial membrane potential phenotypes were also observed in 35 patients with Parkinson’s disease and 25 healthy controls (Payne et al. 2024). An increased mitochondrial and lysosome co-localization and an increased rate of mitochondrial collapse were found in sporadic Parkinson’s fibroblasts, a phenotype reversible by the LRRK2 inhibitor LRRK2-in-1 (Smith et al. 2016). In spinal muscular atrophy, machine learning could distinguish fibroblast images of 12 patients from 12 healthy controls, though the study could not rule out technical confounders (Yang et al. 2019). SORL1 is a gene associated with Alzheimer’s disease; morphology phenotypes distinguished SORL1 knockout neural progenitor cells from isogenic wild-type controls (McDiarmid et al. 2023). 15 people with two genotypes of hereditary spastic paraplegia were distinguished from matched controls using stains for DNA, mitochondria, and acetylated α-tubulin, plus label-free phase contrast cell images in fibroblasts (Wali et al. 2021). Other studies also uncovered phenotypes for neurological conditions in smaller cohorts (Mortiboys et al. 2008, 2010; Blanchet et al. 2015; Smith et al. 2016; Teves et al. 2017; Hung et al. 2018; Wali et al. 2023).

Therefore, although some conditions might only have phenotypes detectable at the organ or organismal level, others do seem to manifest at the peripheral cell level. Similarly, although some disease-related phenotypes might only be visible in a “relevant” cell type, others may be detectable in a broader spectrum of cell types, including fibroblasts. This may be due to the repurposing of many cellular components across diverse functions in different cell type contexts, or because cellular mechanisms may exist and be impacted in a diversity of cell types even though the symptoms experienced by a patient are only observed in a particular organ system, perhaps due to compensatory mechanisms elsewhere. For psychotic disorders, it is notable that patients are at elevated risk for other non-psychiatric disorders and there are substantial overlaps between gene sets associated with the two (Andreassen et al. 2013; Bulik-Sullivan et al. 2015). Therefore, it would not be surprising if peripheral cells showed abnormalities relevant to those in brain cells, consistent with the observation that schizophrenia and bipolar disorder have known associations with skin abnormalities (McPhie et al. 2021). Because obtaining live cells from the human brain is rarely possible and because generating relevant neural cell types by either differentiation of hiPSC lines or by direct conversion of somatic cells is challenging at scale (Panchision 2016), identifying phenotypes in cell types that are readily obtained and high-throughput-compatible could accelerate discovery.

In 2010, we reported finding a mitochondrial localization phenotype in fibroblasts and lymphocytes from 10 bipolar patients as compared to 10 control patients (Cataldo et al. 2010). Specifically, the distribution of mitochondria was unusual in bipolar patients, whose cells displayed mitochondrial networks that were much more dense and concentrated near the nucleus, versus controls whose mitochondria dispersed more evenly throughout the cell.

The finding of image-based abnormalities in mitochondria of patients with conditions producing psychosis is now affirmed by a wide variety of genetic, brain imaging, and peripheral cell studies (Flippo and Strack 2017; Scaini et al. 2021; Papageorgiou and Filiou 2024). Postmortem brains of bipolar patients have more, but smaller, mitochondria (Cataldo et al. 2010), as do their fibroblasts *ex vivo* (Marques et al. 2021) and iPSC-derived neurons, *in vitro* (Mertens et al. 2015). Mitochondrial fragmentation is also seen in Epstein-Barr virus (EBV)-transformed lymphocytes from schizophrenic patients (Rosenfeld et al. 2011). Similar glutamate-induced fragmentation can be mitigated in mouse hippocampal HT22 cells when the psychiatric disorder-associated gene CACNA1C (Sklar et al. 2008; Moon et al. 2018) is knocked down by RNA interference (Michels et al. 2018). Postmortem brains of patients with schizophrenia show a lower density of mitochondria in axon terminals, compared to healthy brains (Roberts 2021). Evidence for functional abnormalities in mitochondria is even more solidly associated with these conditions, ranging from genetics (Cuperfain et al. 2018), alterations in mitochondrial fusion/fission proteins (Scaini et al. 2017), bioenergetics (Zanfardino et al. 2021), mitochondrial membrane potential (Munakata et al. 2004; Robicsek et al. 2013; Brennand et al. 2015; Marques et al. 2021), and more (Scaini et al. 2016, 2021; Pereira et al. 2018; Wu et al. 2019; Kuperberg et al. 2021; Giménez-Palomo et al. 2021). Likewise there is evidence for morphological and functional impairment of mitochondria in major depressive disorder (Culmsee et al. 2018; Chen et al. 2024). Collectively, these findings are consistent with the more general conclusion that mitochondrial morphology/localization and function are tightly related and abnormal in many psychiatric conditions (Bulthuis et al. 2019; Chan 2020; Jenkins et al. 2024; Ni et al. 2024).

Here, we sought to increase confidence in a mitochondrial localization phenotype in psychiatric conditions by testing a new, larger cohort: 168 people as compared to our original set of 20. Indeed, the lack of attempts to replicate findings in larger cohorts has been described as limiting progress in the field (Harrison et al. 2020). We also sought to expand the study to related psychiatric conditions beyond bipolar disorder - for the past fifteen years, we have been collecting samples from patients experiencing psychosis across a spectrum of conditions including schizophrenia, schizoaffective disorder and bipolar disorder, along with healthy control patients and patients with major depressive disorder. Finally, because computational analysis of images has advanced dramatically in the past decade and public datasets are now available, we were also able to develop and pilot a virtual screening approach based on mitochondrial localization that might identify chemical and genetic perturbations relevant to the psychiatric conditions in other datasets.

## Results

### Mitochondrial localization is disrupted in fibroblasts from patients experiencing psychosis

We sought to identify an image-based phenotype in a cohort of patients experiencing psychosis. Success could then power screening public image datasets to identify potential therapeutics by identifying those that impact the phenotype (Figure 1). We collected skin punch biopsies from 168 individuals and derived human fibroblasts from them, including from 50 people with a primary diagnosis of bipolar disorder (BP), 38 with schizophrenia (SZ), 11 with schizoaffective disorder (SZA), 1 with psychotic disorder not otherwise specified, 20 with major depressive disorder (MDD), and 48 healthy controls (HC); together with patient demographic metadata including their primary diagnosis. The median and range of ages was consistent across patient categories, as was the typical male-skewed ratio seen for these conditions, except for MDD (Supplementary Table 1 and Supplementary Figure 1). We grew the cell lines as consistently as possible prior to plating, fixing, staining for DNA, actin, and mitochondria (using MitoTracker Orange-M7510), and imaging (Figure 2A and Supplementary Figure 2).

**Figure 1:**
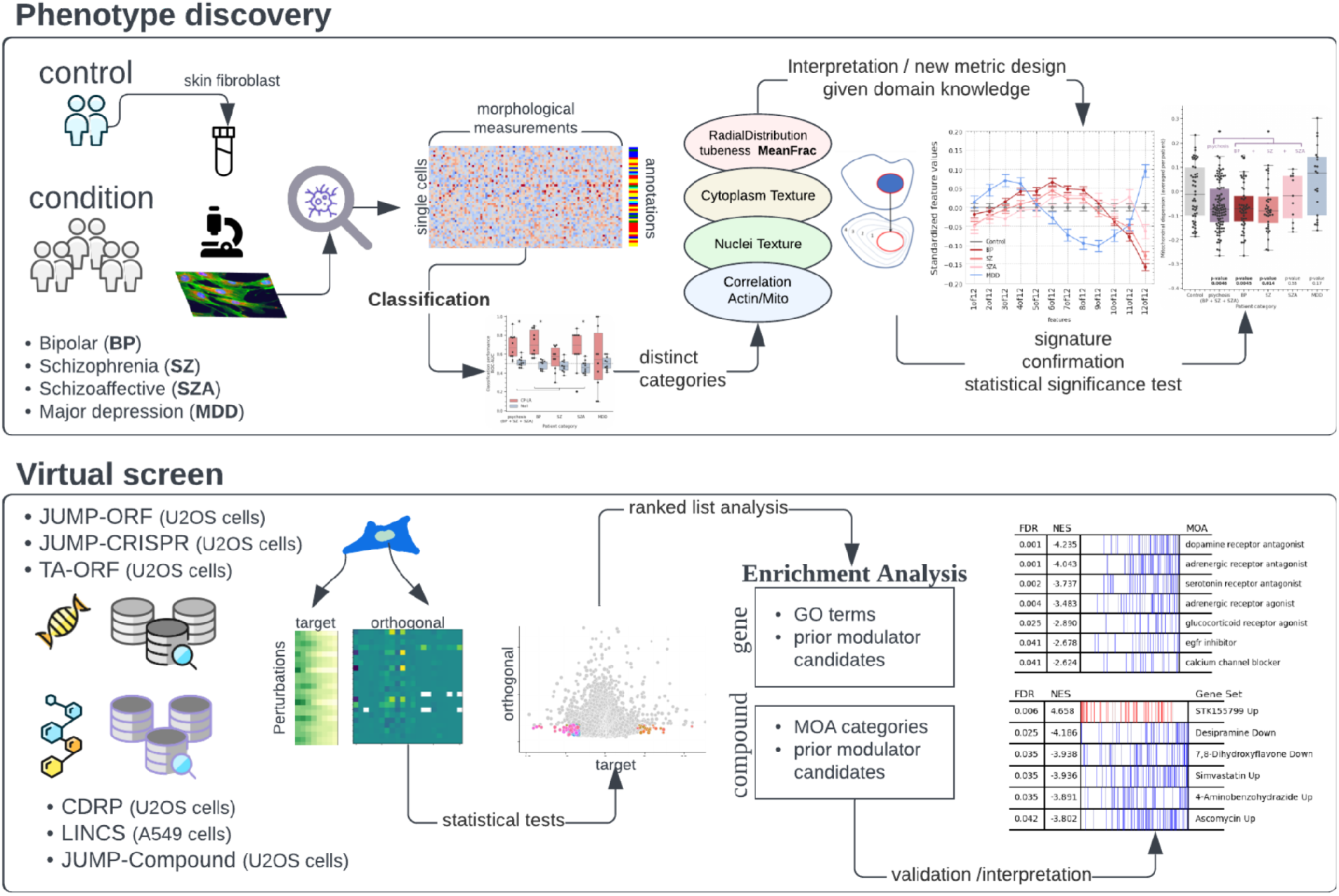
Overview of our strategy for phenotype discovery and virtual screening.

**Figure 2:**
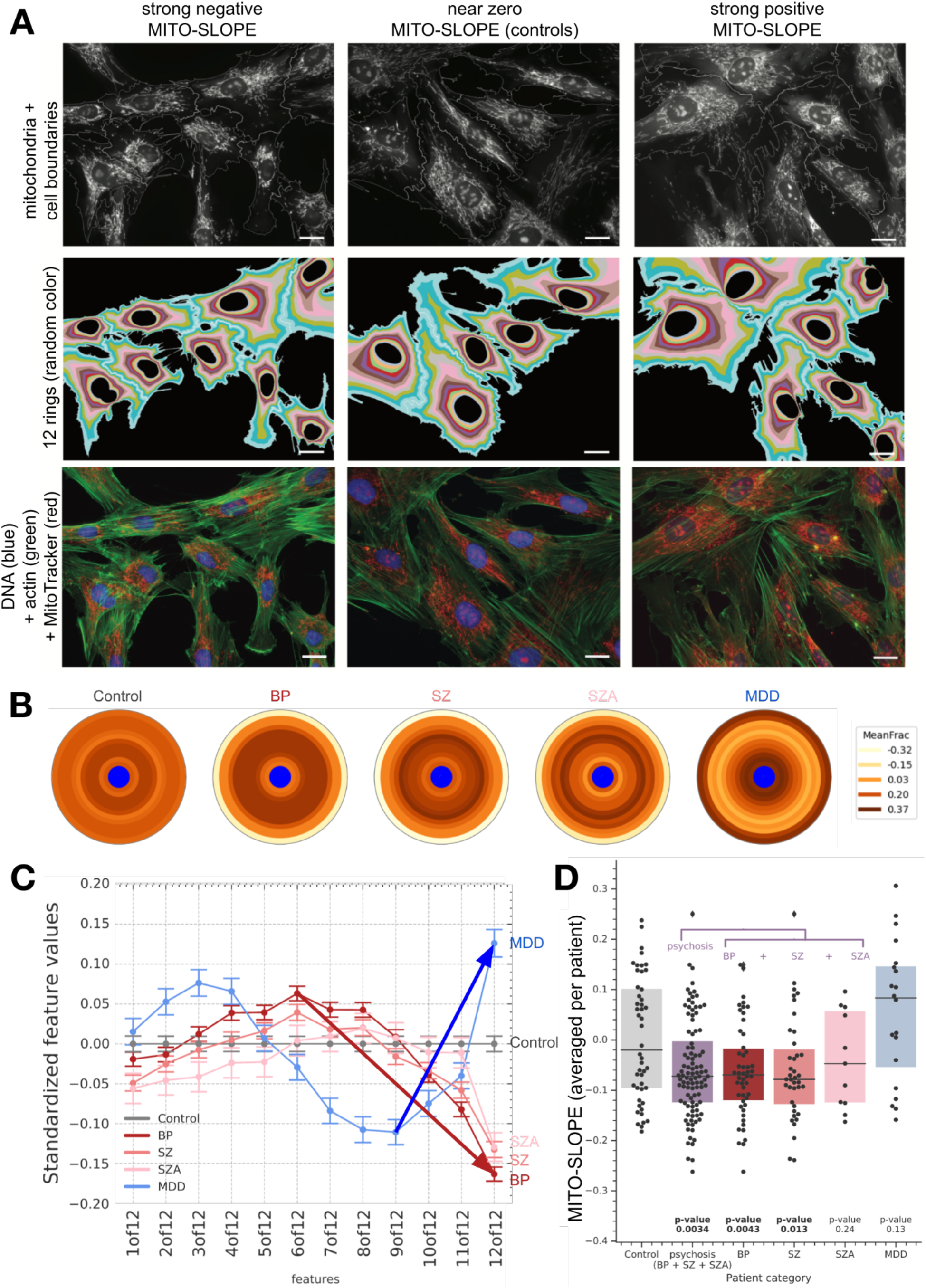
Localization of mitochondrial staining differs in various patient groups experiencing psychiatric conditions. (A) Patient fibroblasts that are stained for DNA, actin, and mitochondria (also shown in grayscale with white outlines for cell border segmentation). Arbitrary colors show the 12 rings used to compute MITO-SLOPE based on segmentation of the nuclear and cell borders, which are imperfect in this high-throughput study. Cells whose nuclei touch the border of the image are excluded from computing MITO-SLOPE. Cells are from subjects with low/median/high slope, but there is significant heterogeneity across cells per subject. Scale bar = 20 μm for all images, which are full fields of view. (B) Localization features computed for MitoTracker staining (after a tubeness transformation, Methods). Integrated intensity of staining is measured in each of 12 concentric rings from the nucleus (visualized in blue) to the cell border, with the shape and size of the rings conforming to the shape and size of the cell (plotted here as if the cell is perfectly circular), and normalized to the fraction of pixels in that ring. Data is z-scored to the control population (for all patient groups, mitochondrial staining lessens towards the cell periphery, so we analyze relative changes). Note that for the Control group, values are not precisely zero because each ring represents a distribution of normalized values that center around 0 coming from a population of individually normalized single cells with biological variability. Patients with bipolar disorder (BP), schizoaffective disorder (SZA) and schizophrenia (SZ) show a declining pattern at the cell periphery, indicating reduced mitochondrial density near the cell edge. In contrast, major depressive disorder (MDD) does not show decreased peripheral mitochondrial signal. (C) Mean standardized values (± standard error) across the 12 rings, ordered from the edge of the nucleus (1) to the cell periphery (12). To simplify to a single feature, we defined MITO-SLOPE as the slope of change for this localization metric towards the cell periphery, starting from the ring with the maximum absolute value (see Methods for definition) (arrows). (D) The value of the MITO-SLOPE feature representing mitochondrial dispersion is shown for each patient in each category. “Psychosis” includes the patients in the categories BP, SZ, and SZA. Box plots show the median and upper and lower quartiles; values exceeding 1.5 x IQR are plotted as diamonds. P-values are noted for each comparison to the control group.

By eye, we observed a dispersed mitochondrial network composed of thin filaments extending to the cell periphery in control patients, whereas mitochondrial staining in patients in the categories experiencing psychosis tended to drop off more dramatically towards the cell periphery. The pattern is subtle and heterogeneous across a cell population. We next sought metrics to quantify these observations objectively, consistently, and efficiently, hoping to use the resulting phenotype to screen existing public image sets.

Many metrics exist to characterize mitochondrial networks in different ways (Leonard et al. 2015; Harwig et al. 2018; Hemel et al. 2021; Chu et al. 2022). Some rely on segmentation of the boundaries of individual mitochondria, to measure properties such as area, aspect ratio, and circularity, as well as their position relative to the nucleus, cell edge, or each other. We required a metric not dependent on segmentation because in our patient fibroblast images most of the mitochondrial mass is organized in more filamentous and interconnected fashion rather than as distinct segmentable objects (Supplementary Figure 3). Segmentation is also infeasible on low-resolution public datasets, undermining our goal of screening them. To preserve interpretability, we used classical image processing techniques to extract well-understood features of morphology, which include measures of size, shape, intensity and texture of the cell components observed using the various stains (Stirling et al. 2021). And finally, we avoided machine learning because a single feature is more readily translatable across cell types and experimental batches, including existing publicly available datasets. Keeping the number of features low is also valuable to conserve statistical power and avoid overfitting that can occur when measuring thousands of image features for only dozens of samples (Lones 2021).

Based on our prior study (Cataldo et al. 2010), our observations of patient fibroblasts and existing public datasets, and our expertise in image analysis, we designed a metric called MITO-SLOPE (Figure 2A-C, with details in Methods). MITO-SLOPE is a localization metric that quantifies the gradient of change in the distribution of the mitochondrial network near the cell edge. First, segmentation of cells and nuclei enables dividing the cytoplasm into 12 concentric rings whose shape and size adapt to each cell (Figure 2A). MITO-SLOPE finds the ring with the most mitochondrial density and then measures the decline in density from there to the cell edge (Figure 2C), identified by available cytoplasmic stains (which vary across the image sets in this study, e.g., phalloidin 488 for the patient fibroblasts). Mitochondrial density is measured as the mean fraction of MitoTracker after transformation with a filter to increase tubeness (named RadialDistribution_MeanFrac_NNof12 in CellProfiler). For all patient groups, raw mitochondrial staining is higher at the edge of the nucleus and decreases towards the cell edge; normalization to controls reveals differences across patient groups (Figure 2B, C).

In our cohort of patient fibroblasts, MITO-SLOPE is decreased (relative to healthy controls) for patients who experience psychosis (p = 0.0034), particularly the subsets of that group with bipolar disorder (p = 0.0043) and schizophrenia (p = 0.013) (Figure 2D). This means their mitochondria are not strongly localized at cell edges, relative to controls. Patients with depression do not show the same phenotype. These findings are consistent with our prior published results on a sample of 10 bipolar patients (Cataldo et al. 2010) but expand to a larger number and include multiple diagnoses that present with psychosis. It is also consistent with our prior work that found a mitochondrial localization phenotype in patient astrocytes for a schizophrenia-predisposing syndrome associated with 22q11.2 deletion (Tegtmeyer et al. 2024). Unsurprisingly, not all patients experiencing psychosis had decreased MITO-SLOPE relative to control patients. This is expected given genetic, diagnostic, and environmental variability among patients and controls, as well as the fact that dramatic changes in cell phenotypes would likely be associated with death in utero or early and substantial cognitive impairments such as those seen in neurodevelopmental disorders.

We considered alternate metrics and potential confounders (Supplementary Figures 4 and 5). Although metrics of cell size are weakly correlated with MITO-SLOPE (maximum 0.39), these do not distinguish patient categories (Supplementary Figure 4A-E); MITO-SLOPE is relatively independent of cell size because the number of rings is fixed and the size/shape of rings scales to fit each cell (see Figure 2A for examples). The weighted average of mitochondrial intensity in concentric rings also poorly distinguishes patient categories although it is correlated with MITO-SLOPE (Supplementary Figure 5A). Some skeleton-based and alternate ring-based metrics do differentiate, but less powerfully and/or capture the phenotype less specifically than MITO-SLOPE (Supplementary Figure 5B, C, E, F). These include a measure of radial intensity staining that replicated our previous results (Cataldo et al. 2010) (Supplementary Figure 5B). We also evaluated machine learning, including deep learning, an area of our expertise. These strategies leverage hundreds to thousands of features and as noted are not as accurately applied to virtual screening. We trained classifiers that could distinguish the combination of the three groups who experience psychosis (BP, SZ, SZA) from healthy controls using logistic regression (Supplementary Figure 6A). A convolutional neural network based on raw images’ pixels yielded roughly similar results (Supplementary Figure 6B). These approaches confirm a psychosis-associated phenotype, but do not appear to dramatically improve upon MITO-SLOPE.

### Mitochondrial membrane potential is lower in fibroblasts from patients experiencing psychosis

The MitoTracker™ Orange CMTMRos dye that we used to measure location has also been used to measure mitochondrial membrane potential (MMP) based on its total intensity in the cytoplasm of the cell (Yatchenko and Ben-Shachar 2021). In our patient cohort, we observed a decrease in cytoplasmic MitoTracker staining in cell lines associated with psychoses (Supplementary Figure 5D). This is consistent with prior studies demonstrating decreased MMP in bipolar disorder (Munakata et al. 2004; Marques et al. 2021) and in schizophrenia (Robicsek et al. 2013; Brennand et al. 2015).

### Chemical regulators of mitochondrial localization

Large public image datasets are becoming available, enabling an intriguing possibility: chemical regulators of relevant pathways, and potential therapeutics, might be identified by assessing disease-associated phenotypes in existing datasets of compound-treated cells. Virtually screening existing datasets saves significant cost and time relative to setting up and conducting physical experiments, and such screening has proven effective in identifying promising chemical regulators (Rohban et al. 2022) and therapeutic candidates (Cuccarese et al. 2020). There are caveats. The approach might not work well if the available data are from a different cell type than the cell type where a disease phenotype was identified. However, our ability to find a psychosis-associated phenotype in skin cells, for patients experiencing a brain condition, provides a hint that this phenotype may be present in many cell types. A virtual screen could also fail if effective chemicals are missed because they would only reverse a phenotype in patients’ cells that show the phenotype and not show any impact on cells from people not having a psychotic disorder (as is likely the case for most cultured cell lines used in public databases).

Image-based screens, where compounds are physically tested for reversion of a disease-associated phenotype of interest, are very common. However, only recently has image-based *virtual* (or *inferential*) screening been demonstrated (Anon. 2021; Rohban et al. 2022; Neumann et al. 2024; Celik et al. 2024). To our knowledge, a large image-based virtual screen based on a phenotype uncovered in patient fibroblasts has not been attempted. The strategy is analogous to connectivity mapping using transcriptional profiles (Lamb et al. 2006). For example, a recent study reported using post-mortem transcriptional signatures to virtually query the large Connectivity Map gene expression database to identify 35 drugs that target genes associated with the shared risk pathways across BP and SZ, including some that interact with CACNA1D (Grotzinger et al. 2023).

We therefore measured MITO-SLOPE in existing Cell Painting datasets to see whether already-known regulators of psychiatric or neuropsychiatric conditions impact the MITO-SLOPE phenotype, as a first step in the eventual identification of novel regulators. We began with the LINCS dataset cpg0004-lincs (Way et al. 2022) because it includes mostly approved drugs, allowing us to analyze drug categories known to treat certain symptoms for patients with schizophrenia and bipolar disorder. Dr. Rakesh Karmacharya (see Acknowledgements) manually annotated such drugs into various categories based on their mechanisms, and we found that none of the known psychoactive categories we annotated were statistically enriched in either direction in our compound list ranked by phenotype strength (Figure 3, Supplementary Figure 7). This result is consistent with our previous finding that mitochondrial localization in fibroblasts was not impacted by lithium treatment, nor did patients’ lithium use alter mitochondrial distribution in postmortem brain samples (Cataldo et al. 2010). We note that existing drugs typically do not clearly and specifically target a single protein or process, so these annotated categorizations are imperfect. Further, fibroblasts do not likely express high levels of the proteins that are the direct targets of most of these psychiatric drugs, nor are fibroblasts part of circuits equivalent to the neural networks that mediate the clinical effects of these drugs. In addition, the therapeutic effects of drugs used for psychosis and depression develop over weeks of treatment. Nevertheless, these results leave open the possibility that the assay has the potential to identify candidate therapeutics working by a novel mechanism of action that does not clearly map to existing drug categories.

**Figure 3:**
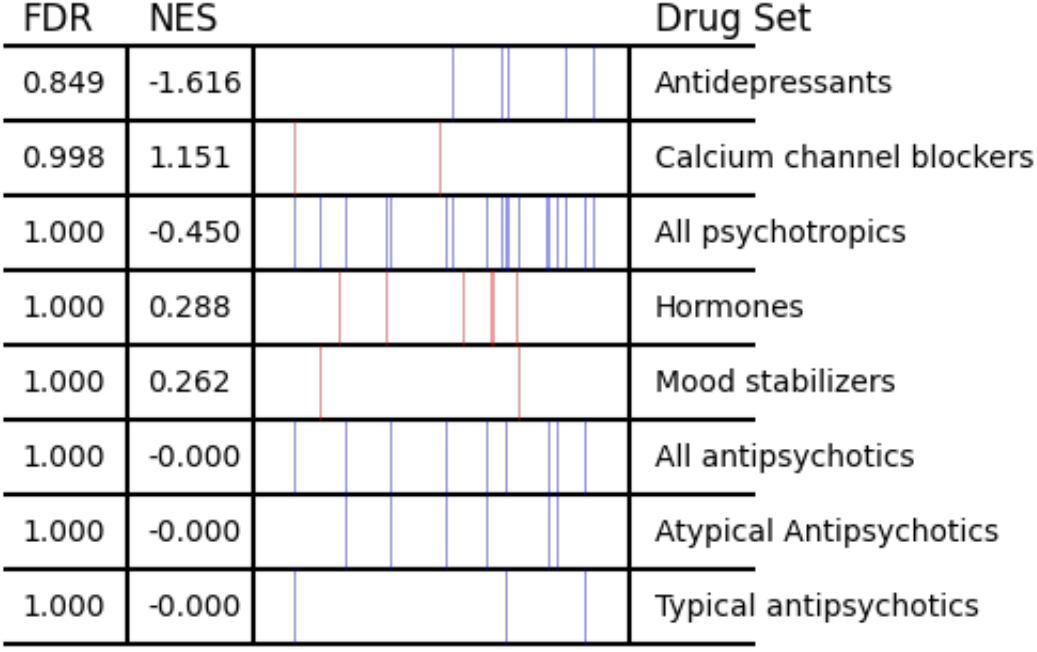
Existing classes of psychoactive drugs do not, overall, show evidence of impacting mitochondrial localization. For drug mechanism-of-action classes with more than four compounds present in cpg0004-lincs, we assessed their impact on the mitochondrial localization phenotype, as measured by the mean value of MITO-SLOPE feature across all cells in the sample. The vertical lines in each row represent the positions of individual compounds from that drug class within the ranked list of all compounds ordered by MITO-SLOPE values. None of the mechanism-of-action classes shows a statistically significant rank ordering (in either direction) relative to the other compounds in the assay. NES: normalized enrichment score (red indicates positive NES: more towards the top of the list of compounds rank-ordered by MITO-SLOPE from positive to negative; blue indicates the class is enriched at the negative end of the list - though, again, none of the categories was statistically significantly enriched); FDR: false discovery rate from Benjamini-Hochberg multiple hypothesis correction across all eight drug classes (see Methods). To avoid loss of signal from samples treated with very low concentrations of each drug, we selected (for each compound) the dose that yielded the strongest impact on this phenotype. Results for all classes of compounds and all doses independently are shown in Supplementary Figure 7. All data were cell-count-corrected.

We then virtually screened the LINCS dataset to find “hits”, drugs that impact MITO-SLOPE strongly in either direction. We quickly noticed that many cancer drugs yield strong changes in MITO-SLOPE but also change many other features of cells that would likely lead to adverse outcomes, and would therefore be undesirable for a drug to treat a chronic mental health condition. We therefore designed a strategy to strictly filter hits to only keep those that minimally change morphology of cells within the intersection of two feature subspaces: (1) features that are orthogonal (do not correlate) to mitochondrial localization, and (2) features that are not distinctive across patient groups in the fibroblast dataset (see Methods).

We found seven positive and three negative MITO-SLOPE hits. The majority are linked to psychiatric or neuropsychiatric conditions (Figure 4), including several drugs currently in active use (citalopram, divalproex-sodium, and tolcapone). Two additional hits, PF-02545920 and trapidil, are known to interact with the phosphodiesterase (PDE) protein family, which has been strongly linked to psychiatric disorders (Hebb and Robertson 2007; Jiang et al. 2023); as noted: “The most promising therapeutic candidates to emerge from these preclinical studies are PDE2 and PDE4 inhibitors for depression and anxiety and PDE1 and PDE10 inhibitors for schizophrenia… PDE3 and 4 inhibitors have shown promising results in clinical trials in patients with depression and schizophrenia” (Sadeghi et al. 2023). Finally, several hits have plausible mechanistic links to psychiatric or neuropsychiatric conditions (pilaralisib, tivozanib, and clioquinol).

**Figure 4:**
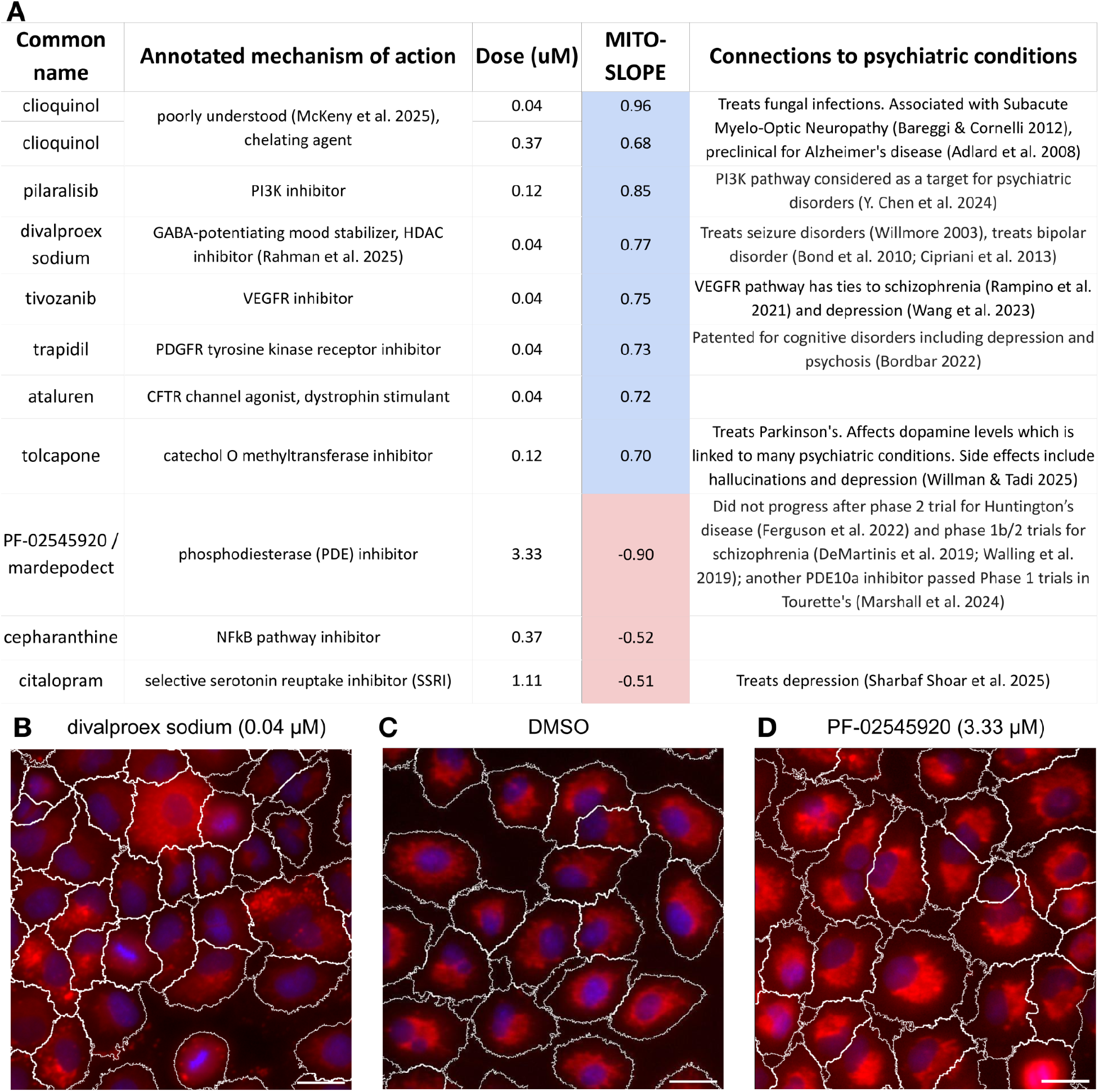
Chemical hits in virtual screen impact mitochondrial localization while minimally impacting cell morphology overall. (A) The table shows compounds that increase (blue) or decrease (red) mitochondria towards the edge of the cells as measured by d_slope (d_slope is the Cohen’s d of the t-test comparing the negative control versus drug-treated cell population, standardized for sample size; the directionality matches the mean of the original values MITO-SLOPE, and the value matches the strength of MITO-SLOPE). Hits were filtered out if they impacted non-mitochondrial-related features (p_orth_std, see Methods). Of note for drug toxicity, the hits do not dramatically impact cell count, within the time frame of the experiment (see Supplementary Table 2, which contains other details and full references). Values were calculated for each compound’s dose independently. Annotated mechanisms are from the Drug Repurposing Hub. (B-D) Representative LINCS dataset images showing A549 cells treated with (B) 0.04 μM divalproex-sodium showing mitochondria spread throughout the cytoplasm, including the cell edge, (C) DMSO vehicle negative control, and (D) 3.33 μM PF-02545920 showing mitochondria condensed towards the nucleus. The LINCS images also contain endoplasmic reticulum, plasma membrane, f-actin, RNA, and Golgi apparatus stains (not shown) that were used during image analysis for segmenting cell borders (white outlines); blue = DNA, red = MitoTracker. Image intensities displayed in the figure were rescaled relatively for each image (using the minimum/maximum of each image - see Methods) for better visualization of the staining pattern but were not rescaled for data analysis. Scale bars = 25 micrometers.

We analyzed MITO-SLOPE in two other public imaging datasets where compounds’ impacts on mitochondrial localization could be discerned and again filtered hits for fairly normal cell morphology based on the orthogonal features. These compound sets have far fewer annotated mechanisms of action because they primarily contain screening compounds rather than well-studied drugs or bioactive agents. In the cpg0016-jump (JUMP) dataset, we found 228 compounds, none with annotated mechanisms of action. In the cpg0012-wawer-bioactivecompoundprofiling (CDRP) dataset, we found 142 compounds as hits. Of these, only two had annotations, both with negative slope and known links to SZ and BP through nicotinic cholinergic receptors. BRD-K28863208 (also known as PNU-282987; MITO-SLOPE = -0.37), is an agonist of the nicotinic cholinergic receptor encoded by *CHRNA7* and is listed as a preclinical program for schizophrenia from Pfizer, terminated for undisclosed reasons (Gray and Roth 2007). The other annotated hit BRD-K85242180 (also known as propyl beta-carboline-3-carboxylate; MITO-SLOPE = -0.56), is an indoleamine 2,3-dioxygenase (IDO) inhibitor: IDO is a key enzyme in the kynurenine metabolic pathway, which has been observed to be dysregulated in both SZ and BP (Erhardt et al. 2017), and kynurenic acid modulates glutamatergic and dopaminergic neurotransmission as well as antagonizing alpha7 nicotinic cholinergic receptors (Erhardt et al. 2017; Plitman et al. 2017). Genetic associations have been observed between alpha nicotinic receptors and both SZ and BP (Thomsen et al. 2011) and smoking nicotine-containing cigarettes is common in both SZ and BP (Heffner et al. 2011; Ding and Hu 2021); nicotine can both reduce and exacerbate mood and cognitive symptoms (Ripoll et al. 2004; Heffner et al. 2011; Ding and Hu 2021).

### Genetic regulators of mitochondrial localization

We next virtually screened our public JUMP Cell Painting images from genetic perturbations, both overexpression (12,609 genes) and CRISPR-Cas9-mediated knockout (7,975 genes) (Chandrasekaran et al. 2024). Because no single gene determines more than a small fraction of risk for SZ or BP, we did not anticipate dramatic impacts on the mitochondrial localization phenotype, but we hoped this analysis might provide insights and further substantiate links between the psychosis-associated mitochondrial localization phenotype and bipolar disorder and schizophrenia.

In the absence of true positive control genes, we took several approaches. First, although not guaranteed to impact the steady-state localization of mitochondria, we analyzed a group of genes annotated as impacting mitochondrial dynamics (fusion, fission, transport, and mitophagy) (Cuperfain et al. 2018). Although there is no guarantee that CRISPR-Cas9-mediated transient knockout was effective for every gene in the set, this group was statistically significantly enriched as lowering MITO-SLOPE (Figure 5A, less mito staining at the edge of the cell, mimicking the psychosis patient samples; Supplementary Figure 8). We did not see enrichment for overexpressed genes in the same categories (Figure 5B), perhaps to be expected given that overexpression of a protein does not guarantee more of that protein’s function and generally produces less robust phenotypes in the JUMP dataset in our previous analysis (Chandrasekaran et al. 2024).

**Figure 5:**
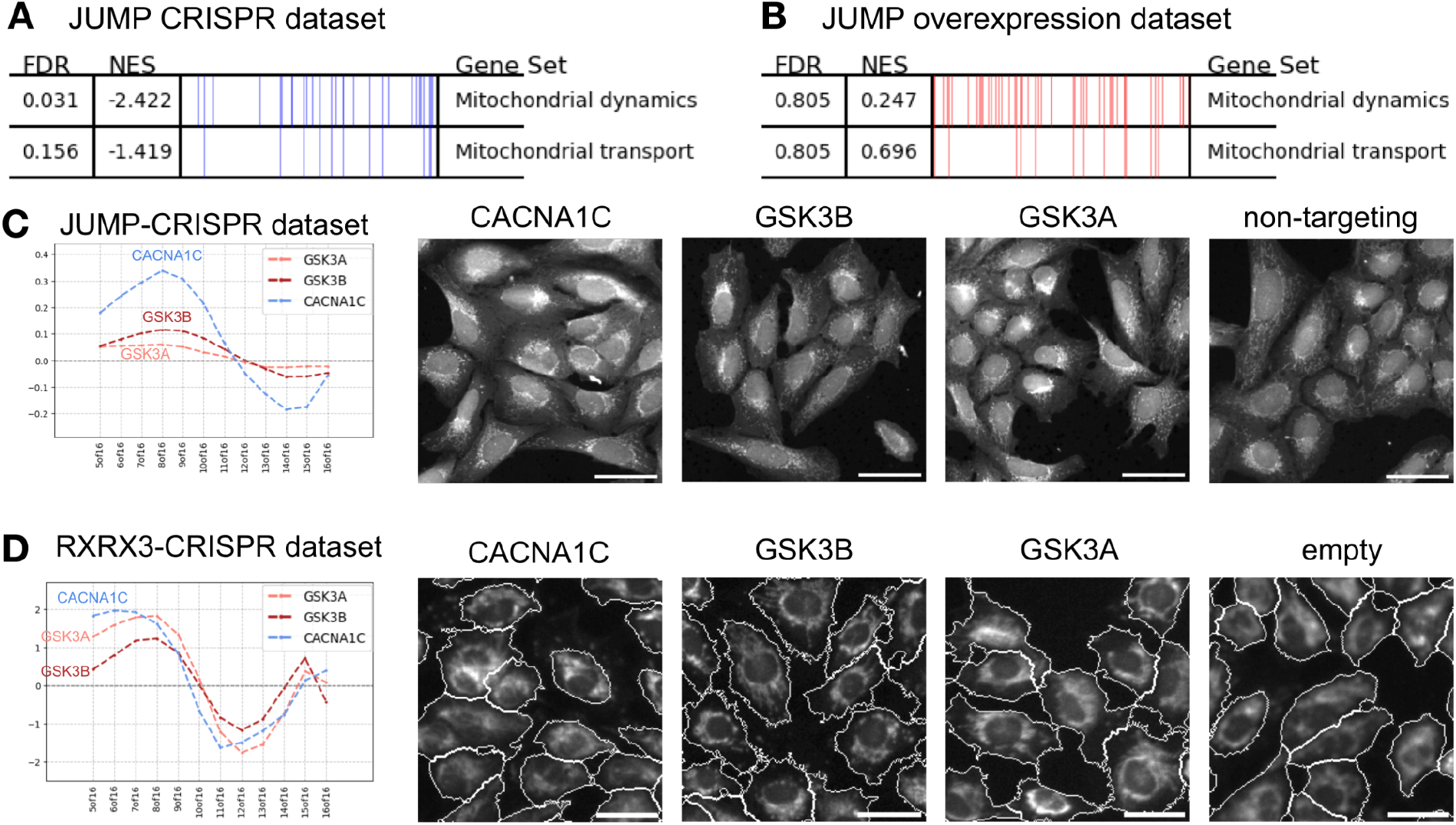
Mitochondrial localization impact for genes with known association with mitochondrial function or psychosis. Mitochondrial dynamic genes are enriched for unusual MITO-SLOPE, for CRISPR knockouts (A) but not overexpressed genes (B); mitochondrial transport genes are not significantly enriched in either dataset. The vertical lines in each row represent the positions of individual genes from that functional category within the ranked list of all genes ordered by MITO-SLOPE values. Gene annotations are taken from Cuperfain, et al., Table 3 (Cuperfain et al. 2018). FDR and NES are as defined in Figure 3. (C and D): As calculated from images in the JUMP-CRISPR dataset of U2-OS cells (C) or RXRX3-CRISPR dataset of Human Umbilical Vein Endothelial Cells - HUVEC (D), we show the distribution of mitochondria in rings from the nucleus (left) to cell edge (right), as in Figure 2C (which shows the patterns of each patient category; all 3 genes here show a pattern more similar to patients with major depression), except this public data used 16 rings, not 12 and the first few rings are omitted because they tend to overlap the nucleus (this dataset’s rings begin at the center of the nucleus, not the edge). Right: Example images from each gene alongside negative controls; an image was selected at random for each condition, cropped to regions with similar cell density representative of the whole image, and scaled similarly for visibility. Both JUMP and RXRX3 datasets contain multiple other channels of stained cellular compartments (shown in Supplementary Figure 8). For RXRX3, the RNA channel was used to segment the cell borders and the resulting cell segmentations are shown. For JUMP-CRISPR, the segmentation pipeline was not provided for the publicly available profiles we used to measure MITO-SLOPE, so segmentation is not shown. Scalebar = 50 μm.

Next, we examined particular genes associated with schizophrenia and/or bipolar disorder: The association of CACNA1C with psychotic and mood disorders is a well-replicated finding (Sklar et al. 2008; Moon et al. 2018) and GSK3B is a known target for bipolar medicines, both antipsychotics (Sutton and Rushlow 2011) and lithium (Snitow et al. 2021), with some variants associated with psychosis in some populations (Jope and Roh 2006) (though, as with most genes, it explains little of risk). We found that CRISPR knockout of CACNA1C had a dramatic pattern, similar to that of major depression, while GSK3A and GSK3B had a much weaker impact (Figure 5C). We analyzed images from a public dataset from Recursion, RXRX3 (Fay et al. 2023), and confirmed the CACNA1C result while also showing a stronger result for both GSK3A and GSK3B (Figure 5D). GSK3B has a well-known role in mitochondrial activity (as reviewed in (Yang et al. 2017)), while CACNA1C has emerging evidence (Michels et al. 2018; Theeuwes et al. 2018; Kessi et al. 2023).

Finally, we analyzed the full list of gene perturbations in both JUMP datasets, to identify those that most strongly impacted the psychosis-associated mitochondrial phenotype (Figure 6). Three genes passed our strict filter for their CRISPR-Cas9-mediated knockout impacting MITO-SLOPE without markedly impacting non-mitochondrial-related features: *SP1*, *TBX3*, and *KCNH6* (Figure 6A). SP1 and TBX3 have clear connections to schizophrenia and to each other; KCNH6 is a potassium channel subunit that modulates cardiac repolarization and heart rate, as well as neuronal excitability, with some links to mitochondrial health. Of note, CACNA1C was at the ∼4th percentile but failed our strict filter for impacting non-mitochondrial features (which could be loosened in reanalysis). Most of the seven genes just below the threshold of significance (RHOU, POLE3, DRD1, KCNA1, FARS2, FOXQ1, ASCL1) have roles in mitochondrial function or psychiatric conditions as well.

**Figure 6:**
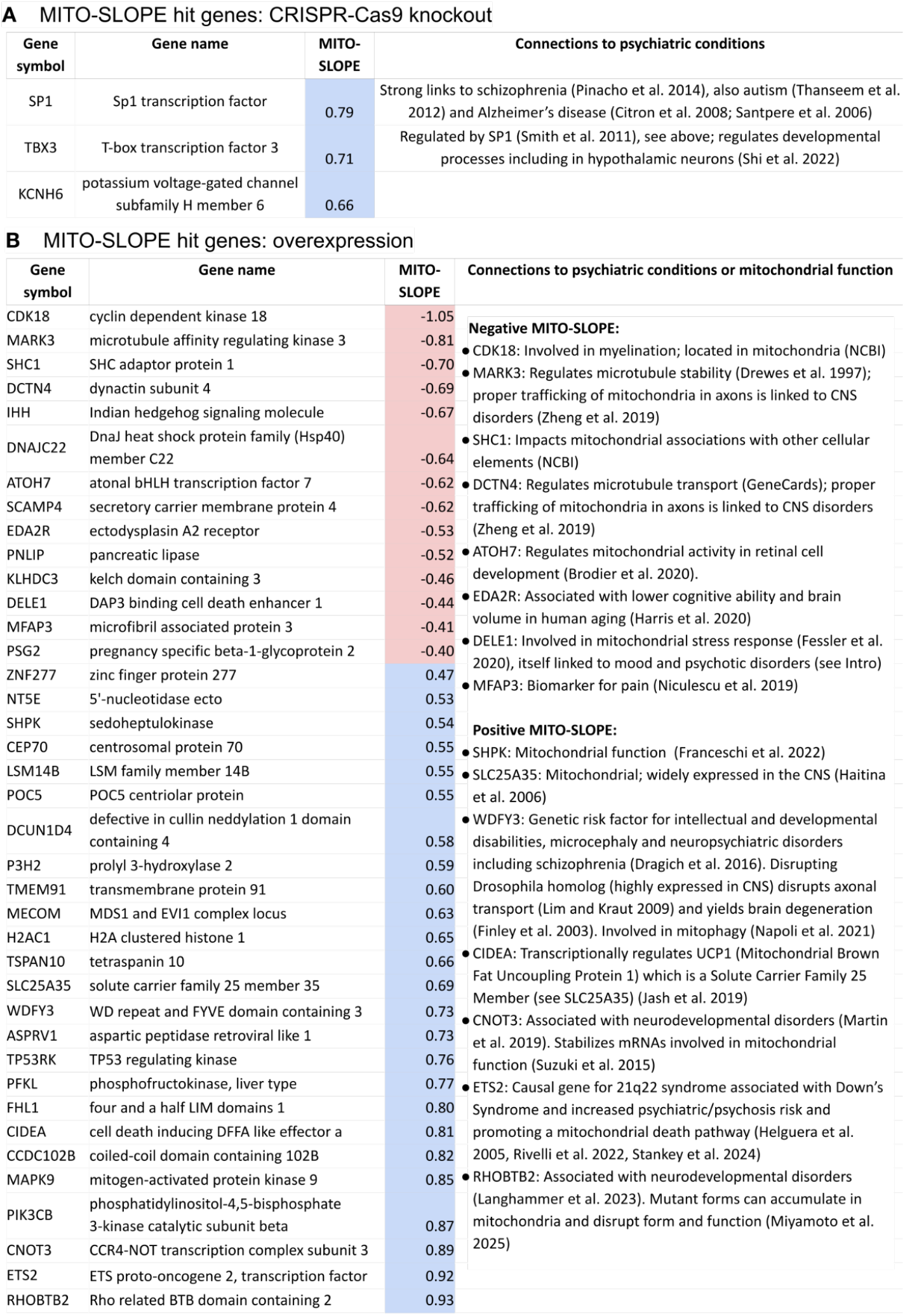
Exploration of genetic hits in virtual screen. The tables show genes that increase (blue) or decrease (red) mitochondria towards the edge of the cells as measured by d_slope and filtered for normal cell morphology (as described in Figure 4 and Methods) in the two datasets (A) JUMP-CRISPR and (B) JUMP-overexpression. The hits do not dramatically impact cell count (Supplementary Table 2, which contains other details and full references).

We found 39 hits in the overexpression data (Figure 6B); enrichment analysis failed to find consistent known annotations among them (Supplementary Figure 9). Nevertheless, among the 14 negative slope hits, we found five that are clearly linked to psychiatric conditions (MARK3, DCTN4, EDA2R, DELE1, and MFAP3) and three related to mitochondria (CDK18, SHC1, and ATOH7). Among the 25 positive-slope JUMP-ORF genes are four genes related to psychiatric conditions (WDFY3, CNOT3, ETS2, RHOBTB2) and three related to mitochondrial function (SHPK, SLC25A35, and CIDEA).

## Discussion

As reviewed in the introduction, the connection between mitochondrial structure/function and psychotic disorders is well-established from multiple streams of evidence. Here, we found in a cohort of 168 individuals that those experiencing psychosis showed mitochondria less localized at the edge of fibroblasts. By querying existing large chemical and genetic perturbation datasets in cancer cell lines, we found that many genes and compounds that impact mitochondrial localization are implicated in brain function and psychiatric conditions.

Transport of mitochondria is tightly regulated and linked to human disorders (Mandal and Drerup 2019). Perinuclear mitochondrial clustering is relatively poorly understood and may be associated with several disorders (Al-Mehdi et al. 2012). The aberrant transport of mitochondria has been linked to schizophrenia and bipolar disorder via several lines of evidence (Flippo and Strack 2017; Scaini et al. 2021). For example, the gene DISC1, Disrupted in Schizophrenia 1, is linked to schizophrenia, depression, bipolar disorder and autism (Murphy and Millar 2017), and mitochondrial dynamics in neurons were impacted by altering DISC1 levels (Atkin et al. 2011) and by large aggregates of DISC1 (Atkin et al. 2012). Mood stabilizers were found to impact proteins involved in mitochondrial transport in synaptoneurosomal preparations from rat pre-frontal cortex (Corena-McLeod et al. 2013). The *C. elegans* protein ANC-1, homolog of human genes (SYNE1/2) associated with various neuropsychiatric conditions, was found to promote localization of mitochondria to the base of the proximal axon (Fischer et al. 2022). Knockout of the schizophrenia-associated protein dysbindin yielded a significant decrease in mitochondrial density in distal axons, which impaired calcium dynamics (Suh et al. 2021). Together with our prior findings including autopsied brain (Cataldo et al. 2010), it appears that the mitochondrial localization phenotype we observe in human patient fibroblasts here is also found in other brain-relevant cell types and in physiological structures *in vivo*.

We can only speculate as to the mechanism by which mitochondrial localization might impact neuron function and ultimately brain function. A recent study found that rather than producing ATP as is typical in most contexts, axonal mitochondria in cortical pyramidal neurons in mice actually consume ATP (Hirabayashi et al. 2024); this builds upon prior work showing that presynaptic release sites lacking mitochondria have a higher likelihood of neurotransmitter release than those with mitochondria (Kwon et al. 2016; Vaccaro et al. 2017). Our findings may be consistent with this: the lack of mitochondrial transport far from the nucleus that we observed here would imply that in psychosis, fewer mitochondria result in more ATP/energy being available in axons, reducing barriers to neurons firing. However, mitochondria produce numerous substrates necessary for cell function, including neurotransmission. These substrates may be lacking in terminals with lower levels of mitochondria. As well, changes in cytoskeletal structure, often involving microtubules, are well-documented in various cell types of patients with bipolar disorder and schizophrenia (Brown et al. 2014; Benítez-King et al. 2016; Marchisella et al. 2016; Snelleksz and Dean 2021). Because some genes associated with bipolar disorders and schizophrenias are also associated with the microtubular cytoskeleton (Perrottelli et al.), drugs targeting this cytoskeleton have been proposed as potentially therapeutic for psychotic and other brain disorders (Varidaki et al. 2018). These processes may also impact mitochondrial localization.

Our patient fibroblast study has several important limitations. First, because of the challenges of obtaining consented patient samples and growing multiple patient cell samples, our cohort is limited to 168, spread across the conditions tested, and was collected in one clinical location with subjective diagnostic annotation. Still, this is larger than most cohorts published for other disorders and offers some mitigation of concerns about donor variability due to factors such as genetics, sex, age, and comorbidities. Second, it is virtually impossible to obtain samples from patients who are untreated with medications, and most patients are on multiple medicines of varying mechanisms. Although the phenotypes we observe are those that persist in cells cultured through several passages and media changes and across patients taking different classes of medicines, we cannot rule out impacts by drugs the patients had been taking. Likewise, we cannot control for other unrecorded factors associated with these conditions, such as diet and exercise. Third, the study assessed a single cell type from patients, fibroblasts, because they are readily obtainable and robust in culture. As noted in the introduction, this strategy has been successful for other brain disorders, and other transport mechanisms have been found consistent across many cell types (Goering et al. 2023). Nevertheless, we have begun to explore mitochondrial localization in more physiologically relevant brain cells, specifically astrocytes derived from the fibroblasts used in the studies reported here - preliminary results show altered mitochondrial localization consistent with that reported in this study; such reprogramming of the fibroblasts to induced pluripotent stem cells (iPSC) followed by induced differentiation to astrocytes is time-consuming but also has the advantage of resetting most epigenetic marks, thus reducing concerns about medication exposure or other environmental differences between patients and controls. Fourth, we report our observations but do not explore the mechanisms underlying the changes in MITO-SLOPE observed: MITO-SLOPE captures mitochondrial localization, which is itself influenced by the intensity of MitoTracker staining, which is influenced by mitochondrial membrane potential - it therefore combines spatial and intensity information, such that disentangling localization from mitochondrial membrane potential is not completely possible. In fact, we believe that the patient phenotype likely reflects multiple cellular changes beyond mitochondrial distribution, including cytoskeletal alterations documented in prior studies. Antibody-based staining for mitochondrial components could provide clarity. And fifth, we performed virtual screening only; no experimental validation of chemical or genetic hits was conducted in patient-derived cells.

Finally, the use of this mitochondrial phenotype as a platform for drug discovery should be approached with caution, as we have previously noted more generally (Chandrasekaran et al. 2020). Most screening strategies that translate findings from cell culture to the human organism are subject to caution; reversing disease-associated phenotypes poses some additional pitfalls. For instance, disease-associated phenotypes may reflect correlation with the patients’ problematic symptoms and not causation. In such cases, drugs reversing the phenotype may be ineffective or “even worse, the detected phenotype might reflect the cells’ attempt to mitigate the impact of the disease perturbation, such that drugs reversing the phenotype would aggravate the condition in patients” (Chandrasekaran et al. 2020). That is, the observed phenotype may be a useful compensation for the abnormalities underlying the symptoms. In that case, reversing the compensation would be anti-therapeutic, although given the tentative observation here of a spectrum of localization with psychosis on one end and depression on the other, medicines at either end might prove useful.

We found that several existing classes of medications used to treat psychosis were not associated with mitochondrial mislocalization. Optimistically, the mitochondrial localization phenotype could represent a novel mechanism by which psychosis might be remedied and is therefore worthy of future chemical screening. Still, there are several reasons why drugs that treat these conditions may not reverse the mitochondrial phenotype in patient fibroblasts nor cancer cell lines. Potential alternative explanations include: (a) the cell type (whether the cancer cell line used in public datasets stained for mitochondria studied here, or a particular patient’s fibroblasts that might be used in a future screen) might be missing key components necessary for proper response, especially the targets of the given drugs. Given that drugs do not work equally well across patients, we would expect heterogeneity in assay response as well, so a future screen would ideally test multiple patient cell lines. As noted earlier, testing more brain-relevant cell types would be ideal, albeit costly, though this may not be necessary for a primary drug screen for compounds working by this mitochondrial localization mechanism. (b) The timing may not be appropriate; existing psychiatric medications typically take 2-3 weeks to show relief of symptoms in patients, although it is also possible that this is only the case due to the complexity of full organ systems. (c) Drug metabolic processes that are only functional in the whole organism may be necessary for the impact of effective drugs in this context. (d) Drug dose, although tested across a range, may be unsuitable.

Future studies could directly evaluate the hit genes and compounds we identified to confirm mitochondrial phenotype reversal and evaluate generalizability in a collection of patient cell lines. Many experimental conditions could be explored for the patient cohort, controls, and perturbations thereof, such as (a) multiple types of cells, such as fibroblasts or differentiated iPSC-derived neural cells (including methylation profiling to capture epigenetic state at biopsy vs. after passaging), (b) multiple readouts beyond MITO-SLOPE, including mitochondrial and cytoskeletal structure and function, mitochondrial transport speed and pausing frequency, membrane potential, and integrity of the microtubular network, and (c) dose-responses of compounds.

It could also be valuable, albeit expensive, to physically screen a broad assortment of compounds or genetic perturbations to find those impacting mitochondrial localization directly in fibroblasts from patients experiencing psychosis rather than this study’s virtual query in unaffected patients’ cancer cell lines. In such a setting, where images are collected in a single well-controlled experiment, searching for compounds that reverse a more sensitive and specific machine-learning based disease phenotype is more feasible, rather than the single feature used here to facilitate virtual screening using publicly available data. On the computational side, more lenient hit-calling in the public image datasets could yield further targets for exploration, and hits could be used to train machine learning models to predict novel neurological targets or to predict mitochondrial localization from chemical structures.

Overall, we found here that the mitochondrial phenotype we originally observed in a small cohort of bipolar patient fibroblasts, and which was seen in our prior study of post mortem brain, is now observed across a much larger group of patients experiencing the most common conditions associated with psychosis, although not in every patient. We find this phenotype can be modulated by genetic and chemical perturbations with plausible links to these conditions, although individual hits require experimental validation. Given the evidence in the literature for mitochondrial function disruption in conditions involving psychosis, and given that the phenotype could be simplified to a single feature, the mitochondrial localization phenotype may be a useful readout for further virtual and physical screening to identify novel targets and therapeutic candidates for those patients whose cells show this phenotype.

## Methods

The protocol for cell culture, sample preparation, and imaging is described in Methods but also provided online (https://github.com/carpenter-singh-lab/2025_Haghighi_Mito/blob/main/McleanCollectionFibroblastGrowthProtocol.md).

### Primary cell lines

Fibroblasts were derived as in (McPhie et al. 2018). All subjects provided written informed consent, as approved by the McLean Hospital/Mass General Brigham (formerly Partners Healthcare) Institutional Review Board IRB# 2007P002597.

Human fibroblast cell lines were grown as in our prior study (Cataldo et al. 2010). Three passages were done prior to plating onto glass coverslips for staining and imaging. Cells were plated at a density of 7.9 x 10^3^ cells per cm^2^. MitoTracker-M7510, 125 nmol/L (Invitrogen Corporation, Carlsbad, CA), was used to visualize mitochondria according to manufacturer’s instructions, with a 30 minute live incubation before fixation and two wash steps prior to fixation with fresh paraformaldehyde. Cytoskeletal integrity was examined by immunocytochemistry using Phalloidin-488 (1:500) according to the manufacturer’s protocol and nuclei were visualized by Hoechst 33342 trihydrochloride (2 ug/ml). Two coverslips per line were mounted in Fluoro-Gel aqueous mounting media (Electron Microscopy Sciences, Hatfield, PA).

### Imaging

For each patient sample, fifty random fields were selected manually and imaged at 400X in three independent channels on a Zeiss Axiovert Observer 2.1 with a Colibri 2 LED illumination system. Three research assistants acquiring images of the samples were blind to the subject group of the sample. The first field was chosen and focus offsets determined for each of the two coverslips on a microscope slide. All of the other fields per coverslip were acquired by moving the X Y dials on the microscope stage. The operator adjusted the focus in the mitochondrial channel on the computer monitor if needed for the sharpest focus prior to acquiring the image. Image acquisition was done with Zeiss Zen 2.0 software using consistent imaging settings from condition to condition, including exposure times - for example, channel settings: DAPI 40 ms/40%, brightness, offset -0.80 um. Green 400 ms/50% brightness. Red 800 ms/100% brightness. Single channel images were exported as high resolution TIFF files as both color and black and white images as well as one merged three-color image. Images were visually checked for focus, defined edges on each cellular element to be analyzed. Cells with nuclei touching the border of the image are excluded from analysis by the automated pipeline. Image fields of view were selected to avoid regions where cells significantly overlapped/grew on top of each other. Images had 8.5 +- 2.2 (median +- stdev) cells in them showing that they had a consistent number of cells. Overall, segmentation quality was good, for primary fibroblasts.

For visualization in Supplementary Figure 2, each fluorescence channel is independently normalized by its maximum intensity and contrast-enhanced by clipping to the 0.5th–99.5th percentiles, then rescaled to 8-bit (0–255). The composite image is generated by averaging the colorized channels and rescaling to 0–255. The final column shows grayscale crops from the mitochondrial channel after per-image max normalization but without additional percentile clipping. All intensity adjustments are applied for visualization only, as absolute staining intensity is not the phenotype of interest in this figure.

### Image analysis and feature engineering

A set of images, obtained as described above, of nuclei, cytoskeleton, and mitochondria were processed by an image analysis pipeline to quantify the mitochondria within individual cells using CellProfiler 3 (McQuin et al. 2018) or CellProfiler 4 (Stirling et al. 2021). The nucleus was segmented using a seed-based watershed algorithm on DAPI images. The cell objects were seeded from nuclei objects and were segmented on actin images or on mitochondria images and then expanded to actin images. We also applied filters to remove cells touching the border, with segmentation issues, and with intensity artifacts as described in the Github repository for the project. The CellProfiler pipelines for processing fibroblast data are available at the project’s github repo and contain detailed parameters; here we describe each step of this pipeline for the patient fibroblast cohort:

1) **LoadData:** Single-channel grayscale images are loaded for DAPI (Hoechst), Phalloidin488 and MitoTracker (m7510) channels. Relevant Metadata and Groups are set.
2) **IdentifyPrimaryObjects**: This step segments the stained nuclei. To aid correct segmentation of the nuclei as objects, objects are excluded based on extreme size, and based on touching the edge of the image. An automatic declumping algorithm is applied in this step to separate adjacent nuclei.
3) **IdentifySecondaryObjects:** A two-step strategy is used to identify the cell edge. First, the mitochondrial network image is used to get close to the outer edge of the cell and this is called the “precell”. Classical segmentation methods perform best when a stain is bright in the center of the object and change (usually fade) towards the edge; actin stain is too consistent throughout the cell body to be a good stain for classical segmentation in images where cells touch. Using the multi-step method allows for the precell to be a large seed and the actin image is used to get the correct segmentation starting from the edge of the precell all the way to the outer cell edge.
4) **IdentifyTertiaryObjects:** This step segments the “cytoplasm” objects by subtracting the Nuclei object from the Cells object.

A series of measures are then done on the segmented objects: The idea is to measure a large number of parameters that will be used in the unbiased machine learning analysis to determine where the largest differences are seen between the groups.

5) **MeasureColocalization**: This module applies various algorithms measuring co-localization within the segmented objects.
6) **MeasureGranularity:** This module measures granularity within the segmented objects. Granularity is a texture measurement that erodes signal in an image using increasing sizes of erosions and quantifies the amount of signal lost from the image in each subsequent erosion.
7) **MeasureObjectIntensity:** This module measures a number of intensity features for segmented objects.
8-10) **MeasureObjectNeighbors:** These three modules quantify spatial relationships between objects of interest.
11) **MeasureObjectIntensityDistribution:** This module measures the spatial distribution of intensities within an object using concentric annuli (rings) from a designated point. At this point of the pipeline, this step is done considering the Identify Primary, Secondary and Tertiary objects (nuclei, cells, and cytoplasm).
12) **MeasureObjectSizeShape:** This module measures many area and shape features (area, perimeter and others) of segmented objects.
13) **MeasureTexture:** This module measures the roughness or smoothness of the staining in an object by measuring the intensity variations in their grayscale images.
14) **EnhanceOrSuppressFeatures:** This module enhances or suppresses the intensity of certain pixels relative to the rest of the image, by applying image processing filters to the image. It is used in this pipeline to enhance the staining of the mitochondrial network so that the overall spatial distribution of the mitochondrial staining can be identified and then measured in a later step. It uses a tubeness filter (algorithm) that enhances filamentous (neurite like) structures of a specified thickness (https://www.longair.net/edinburgh/imagej/tubeness) (Sato et al. 1998). At the end of this module the tubeness image is intensity-rescaled. In essence, this module generates a compromise between the original intensity image and a segmented image. Its purpose is to sharpen the intensity staining without completely turning it into a binary image.
15) **Threshold:** A threshold filter is then applied to the tubeness image in order to determine what intensities are considered foreground and background; this is used in the next module.
16) **MorphologicalSkeleton:** Creates a skeleton from the thresholded mito tubeness image. It delineates the area of the image to be measured in the next module.
17) **MeasureObjectSkeleton:** This module measures the number of trunks and branches for each branching system in an image. The module takes a skeletonized image of the object plus previously identified seed objects and can generate measures of the complexity of the branching object, in this case the contiguous parts of the mitochondrial network.
18-20) **ImageMath:** This module is typically used to do simple mathematical operations on image intensities. Here, we use it without performing a mathematical operation to simply duplicate an image for renaming to be used in a downstream operation, in order to have three independent image names for the three types of radial distribution measurements in the three MeasureObjectIntensityDistribution modules. This is necessary in this pipeline to prevent naming collisions in measurements output by MeasureObjectIntensityDistribution.
21-23) **MeasureObjectIntensityDistribution:** The mitochondrial pipeline uses three MeasureObjectIntensityDistribution modules. Each measures the radial distribution of the enhanced mito tubeness image in a fixed number of concentric annuli (rings) that are scaled across each cell individually. They differ by where the inner edge (center) of the first annulus is set. We separately measured from the center of the nucleus (CCN), edge of the nucleus (CEN), and center of the cell (C). For the patient fibroblast images, the measurement used in the MITO-SLOPE calculation is based on the inner edge of the first annulus being set at the outer edge of the nucleus (CEN); other datasets use other centers (described below).
23) **MeasureImageIntensity:** This module measures the intensity of the whole image and the intensity of the background, in regions without cells.
24-25) **OverlayOutlines:** This module overlays an outline of an object of interest on an image used in the pipeline. We used it to draw the nucleus and cell object outlines onto a black background so that the exact object outlines identified in the pipeline could be saved.
26-27) **Save images:** At this point in the pipeline this module saves an image of the outlines created in OverlayOutlines step on an earlier image from the pipeline.
28) **ExportToSpreadsheet**: Data (Measurements) are exported as .csv files for further analysis.

For displaying the example hit images for the LINCS dataset, intensities were rescaled relatively, for visualization, as follows: the MitoTracker image is raised to power of 0.4 then rescaled to the min/max for each image. Then, the red-green-blue merged image is made with MitoTracker and DNA as red and blue, respectively, and rescaled with a set range of .05-.7. Finally, cell outlines from CellProfiler are overlaid in white.

### Patient group classification using machine learning

We performed patient category classification for various psychosis spectrum subgroups versus control using the patient-level profiles. Each patient has a bag of single cell samples and we form per-bag representations as the median of single cell samples representations in each bag when training a traditional ML classifier. We used a subset of mitochondria channel features extracted by CellProfiler for learning a logistic regression classifier. We used raw single cell mitochondria channel images for learning a CNN deep learning based classifier and we evaluated performance of each learned model through stratified K-fold cross validation (K=10). The area under the receiver operating characteristic curve (AUC-ROC) is reported for each binary classification as the measure of classification performance.

CellProfiler features extracted from the mitochondria channel form a set of 326 highly correlated features. This multicollinearity in our data could affect the feature importance exploration analysis. We handled multicollinearity in our data by forming subgroups of highly correlated features and selecting one feature from each cluster as the feature selection criteria. Subgroups of features were formed by application of hierarchical clustering on Pearson correlation of features and median of each cluster was selected as the representative feature for each subgroup. Supplementary Figure 10 shows the clustering dendrogram and feature subgroups. Y-axis threshold for forming clusters was selected to form a maximum number of clusters while keeping a Variance Inflation Factor (VIF) less than 10 for each feature in the final set. This resulted in a set of 45 representative features used for feature importance exploratory analysis. We note that while using this representative set degrades the classification performance results, it was a necessary step for feature interpretation analysis which is crucial in this study. Classification results reported in Supplementary Figure 6 consider all of the set of mitochondrial features, however the feature importance analysis was based on the models learned from the representative set of features (Supplementary Figure 11). This analysis examined only the patient categories that had an AUC-ROC > 0.6 even after using the representative set of features for classification. Features with an AUC-ROC< 0.6 were considered to be indistinguishable from the Control class based on this analysis.

### Feature interpretation and design of the MITO-SLOPE metric

In our effort to select a metric for virtual screening, we considered many factors in addition to the power to distinguish patients experiencing psychosis. We wanted the metric to not involve combining multiple features, which requires choices about their relative normalization and balance. We wanted a metric that could be robustly measured across datasets, especially the public datasets we planned to use for screening, which are lower resolution than our patient fibroblast study and had a particular set of pre-measured features. We wanted a metric that was flexible to changes in typical cell sizes and measured features across experiments.

To make this choice, we began by examining the 45 representative features from the feature importance analysis, in the context of our knowledge of image analysis and our visual observations over the years for cells across these psychiatric conditions. This analysis is based on the logistic regression model’s coefficients and permutation importance; the top ten features from this analysis are reported in Supplementary Figure 11. This revealed that the following categories of features had the most distinctive power across various patient categories versus control samples:

a. A subset of texture measures in the nuclei and cytoplasm of the mito channel.
b. A subset of features measuring the integrated intensity of mitochondria in concentric rings across the cell, where the size and shape of the rings conform to the size and shape of the cell. These features capture the spatial patterns of the mitochondrial network across the cell and the potential shifts of this network toward the nucleus center or cell edge.

Based on the goals of our experiment and the properties of the data sets we wanted to analyze, we chose to calculate MITO-SLOPE on a single series of a measured feature in category (b): Cells_RadialDistribution_MeanFrac_mito_tubenessXof12 (CEN), where X ranges from 1 to 12. This feature is measured on the tubeness-filtered image (see “EnhanceOrSuppressFeatures module” in the *Image analysis and feature engineering methods* section)

The RadialDistribution_MeanFrac feature is a measurement of the total intensity of pixels in a given ring, normalized to the fraction of pixels that are in that ring. In other words, it is the mean fractional intensity at a given radius. MeanFrac is then z-scored and normalized to the control population before it is used to calculate MITO-SLOPE (detailed below).

MITO-SLOPE is calculated based on the pattern of change of MeanFrac, making it fundamentally a localization metric that quantifies the gradient of change in mitochondrial network density towards the cell edge. Figure 2B and C shows this pattern from the nucleus edge to the cell edge across 12 rings defined in the cytoplasm area of a cell. The promise of this metric lies in its adaptability to cell size, cell-to-nuclei size ratio, cell segmentation errors, staining intensity, image resolution, and other sources of batch-to-batch and cross-dataset variations. It does not require mitochondrial segmentation and is not impacted by changes that occur to mitochondria upon certain fixation techniques (Whelan and Bell 2015). MITO-SLOPE measures the slope of the change in the normalized intensity values in each ring, focusing on the final rings close to the cell edge. In some public datasets in this study, the rings defined for extracting features either start from the cell center (C for “Cell” for LINCS dataset) or from the nuclei center (CCN for “Cell from Center of Nucleus” for the rest of genetic or chemical datasets) rather than from the nucleus edge (CEN for “Cell from Edge of Nucleus”). Therefore capturing the pattern at the final rings makes the pattern more transferable across datasets used for virtual screening and avoids any impact from nuclear/nucleolar staining caused by MitoTracker, a documented phenomenon (Qi et al. 2018). “Final” here is defined below (“last extremum”).

MITO-SLOPE is negative for psychosis (less mito at edge) so potential drugs should yield positive MITO-SLOPE; potential treatments for depression might have negative MITO-SLOPE.

**Algorithm for calculation of MITO-SLOPE**

*Inputs:*

- A table containing single cell profiles of all the population with mito_tubeness radial-distribution columns for MeanFrac (Cells_RadialDistribution_MeanFrac_mito_tubenessXof12) across all radial rings. Radial rings are the 12 rings ordered from nucleus edge to cell edge (indices: RING-11 … RING).

- Labels for each cell (“Control” vs. patient groups).

*Output:*

- **MITO-SLOPE** (a scalar per subject).

*STEPS:*

**1. Column-wise standardization (z-scoring)**

Standardize each feature so that each column of the table has mean 0 and SD

1. All subsequent steps use these z-scored values.

**2. Per-subject aggregation (MeanFrac)**

For each subject, average the z-scored MeanFrac radial-ring columns across that subject’s cells to obtain a per-subject profile; *I_sub_*_j*ect*_[*i*] for i = 0 … B−1 (B = 12).

**3. Control mean profile**

Compute the mean profile across all Control subjects; *I_control_*_,*mean*_[*i*]

**4. Control differencing**

For each subject, form *ΔI*[*i*] = *I_sub_*_j*ect*_[*i*] − *I_control_*_,*mean*_[*i*]

**5. Smoothing**

If desired (required on datasets with noisy measurements), apply a light 1D smoothing to *ΔI*. (we applied a Savitzky–Golay filter to ΔI using window_length = 5 and polyorder = 3 (length preserved). This reduces noise while retaining local trends.)

**6. Last extremum**

Find the positions of the global maximum and global minimum of *ΔI*. Restrict to indices 0 … B−3 (exclude last 2 rings prone to segmentation errors); if neither extremum lies in this range, set MITO-SLOPE = 0 (no valid last extremum).

Otherwise, take the later (rightmost) of the two extremum positions within 0… B−3 as the “last extremum.”

**7. Slope to the edge**

Compute **MITO-SLOPE** as:

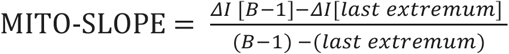

Units: z-score change per ring.

Interpretation:

- MITO-SLOPE > 0 → peripheral enrichment of mitochondrial signal relative to controls.
- MITO-SLOPE < 0 → central shift relative to controls.

### Analysis for virtual screens of chemical and genetic perturbation datasets

We perform the following data processing and analysis on each of the chemical and genetic perturbation datasets to obtain a ranked list of perturbations that modulate the target metric corresponding to mitochondrial dispersion relative to its untreated or control state.

We form a set of features based on patient fibroblast data that have weak linear correlation with the target metric (filtering to less than a 0.3 absolute correlation coefficient with the target metric) and are also not distinctive across various groups of patients versus the control group (filtering to less than a certain threshold defined on the mutual information between the features and the class labels). We decrease redundancy in the set by reducing the features so that those in the filtered set have less than a 0.8 correlation coefficient with each other. We call this the *orthogonal* set of features and use it as a control set of features in which minimal variability is desired. This means we do not want the genetic or chemical perturbations to dramatically change the other dimensions of cell morphology while modulating MITO-SLOPE. The threshold p-value is set based on multiple hypothesis-corrected (adjusted) p-values that therefore differ across datasets. The values of target_bh_corrected_critical_dict were:

{’taorf’: 0.00409,

’lincs’: 0.00761,

’CDRP’: 0.00924,

’jump_orf’: 0.00198,

’jump_crispr’: 0.007,

’jump_compound’: 0.00365}

Note that while the extracted image feature sets are almost consistent across various chemical and genetic datasets, the set of features and channels in patient fibroblast data differ from those in the perturbation datasets. We derive the orthogonal set based on the fibroblast dataset and based on the overlap of those features with those available in the other datasets. As well, during revision we re-examined all patient annotations and as a result reclassified subject 272 and MCL004; this did not impact conclusions of the paper and given the minimal impact of the reclassification on orthogonal features, we kept the original version of them for simplicity.

Perturbation datasets are described in the next section “Screened datasets”. Each perturbation within a batch exists in multiple wells across a variable number of replicate plates. Each well consists of multiple sites, and per-site profiles are derived by averaging features across all single cells available in that site. The MITO-SLOPE metric is then derived from per-site profiles of the RadialDistribution pattern.

To mitigate the effect of plate-to-plate variation in our statistical analysis, we perform a two-sample t-test on per-site MITO-SLOPE values for control versus each perturbation within each plate. We also compare the orthogonal to the target set of features for control versus each perturbation using Hotelling’s T² test. After forming a set of per-plate t-statistics for each perturbation, we select the plate with the median value of the target metric and retain its statistics as the final phenotype strength properties for that perturbation.

We also derive Cohen’s d value and standardized p-value to account for variable sample sizes when comparing each perturbation versus control samples. The final list of perturbations is ranked by Cohen’s d values and filtered to include only significant differences according to the corrected p-value using Benjamini-Hochberg multiple hypothesis correction.

### Mean Fraction of Total Stain Profile Curve Comparison

The area between the mean profile curves for Cells_RadialDistribution_FracAtD_mito was calculated by dividing the region between curves into subregions of trapezoidal and triangular shape and summing their areas. All areas contributed positively to the sum regardless of the relative order of bipolar and control means. We used 10,000 permutations of the diagnostic labels to approximate the distribution of the area between curves under the null hypothesis of no difference in distribution by diagnosis. A p-value was calculated as the fraction of permutation-based areas between curves greater or equal to the value observed for our data.

### Screened datasets

The chemical and genetic perturbation datasets used for virtual screening have previously been published and described. The cpg number allows freely accessing the images and related data from the Cell Painting Gallery (https://github.com/broadinstitute/cellpainting-gallery). Briefly, the datasets include:

● cpg0004-lincs (LINCS, sometimes called LINCS-Pilot1), which is 1,571 compounds across 6 doses in 5 replicates in A549 cells (Way et al. 2022). Images can be viewed here https://idr.github.io/idr0125-way-cellpainting/
● cpg0012-wawer-bioactivecompoundprofiling (CDRP, also called CDRP-BBBC047-Bray), which is 30,430 compounds in U2OS cells at a single dose each, and in 4 replicates per compound (Wawer et al. 2014; Bray et al. 2017). Images can be browsed at Image Data Resource as idr0016.
● cpg0016-jump (JUMP) which is 116,000+ compounds and 15,000+ genes (CRISPR knockout and overexpression) profiled in U2OS cells (Chandrasekaran et al. 2023, 2024). Subsets of images can be browsed at third party websites mentioned therein.

### Enrichment analysis

We performed an enrichment analysis in Figures 3 and 5. Virtual screen analysis of each of the genetic or chemical datasets determines the strength of the target phenotype for each perturbation with respect to the control population through t-test statistics. We sort this differential image signature list by the Cohen’s d values and inspect enrichment of any candidate gene/compound set towards the top or bottom of the overall set using the GSEA strategy (Subramanian et al. 2005). The degree of a candidate set being overrepresented towards the extremes of the list is evaluated by an enrichment score (ES) which represents the deviation from a uniform distribution of the candidate set throughout the list. To check for statistical significance of a calculated enrichment score, p-values are calculated by forming a null ES distribution and permutation testing. We used blitzGSEA (Lachmann et al. 2022) for the enrichment analysis. Enrichment scores normalized by the size of the gene set (NES) and false discovery rate (FDR) for the presented NES values are reported for each set under enrichment analysis. FDR values are calculated using the Benjamini-Hochberg procedure to correct for multiple hypothesis testing across all gene/compound sets being analyzed.

## Data and Code Availability

Raw fibroblast cell images and single cell image profiles extracted by CellProfiler are available at http://doi.org/10.5281/zenodo.15390513. The CellProfiler pipelines for extracting them and source code to reproduce and build upon the presented results are available at the project’s github repository (https://github.com/carpenter-singh-lab/2025_Haghighi_Mito). Several results files are located in the results subfolder there. We license the source code as BSD 3-Clause, and license the data, results, and figures as CC0 1.0.

## Acknowledgements

We gratefully acknowledge input from other members of the Cohen, Cimini and Carpenter– Singh labs as this project developed. In particular, we thank Dr. Yu Han for running preliminary analyses and Dr. Rakesh Karmacharya for manually annotating drugs into mechanism-of-action categories.

## Supplementary information

**Supplementary Table 1:** A Word document containing subject demographics, comorbid conditions, and medications at time of biopsy. A summary of these data, including diagnosis, sex, and age is provided as Supplementary Figure 1.

**Supplementary Table 2:** An Excel file containing details about the chemical and genetic hits in the virtual screens, with full references.

**Supplementary Figure 1.**
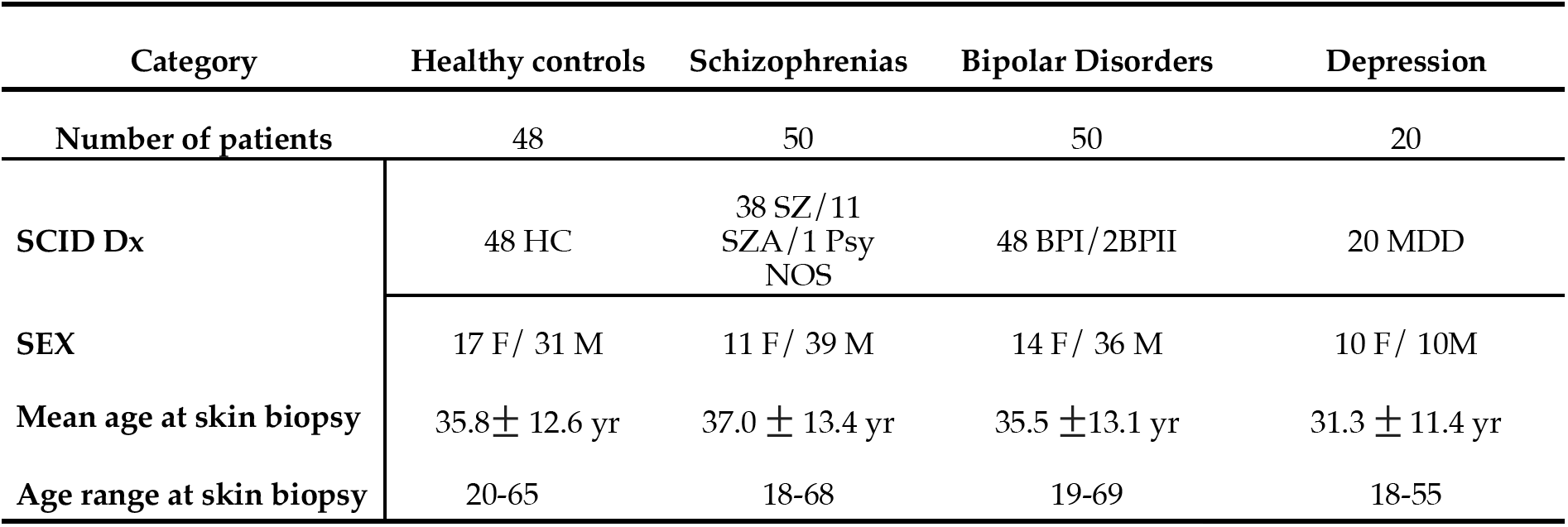
Characteristics of patients in the study.

**Supplementary Figure 2.**
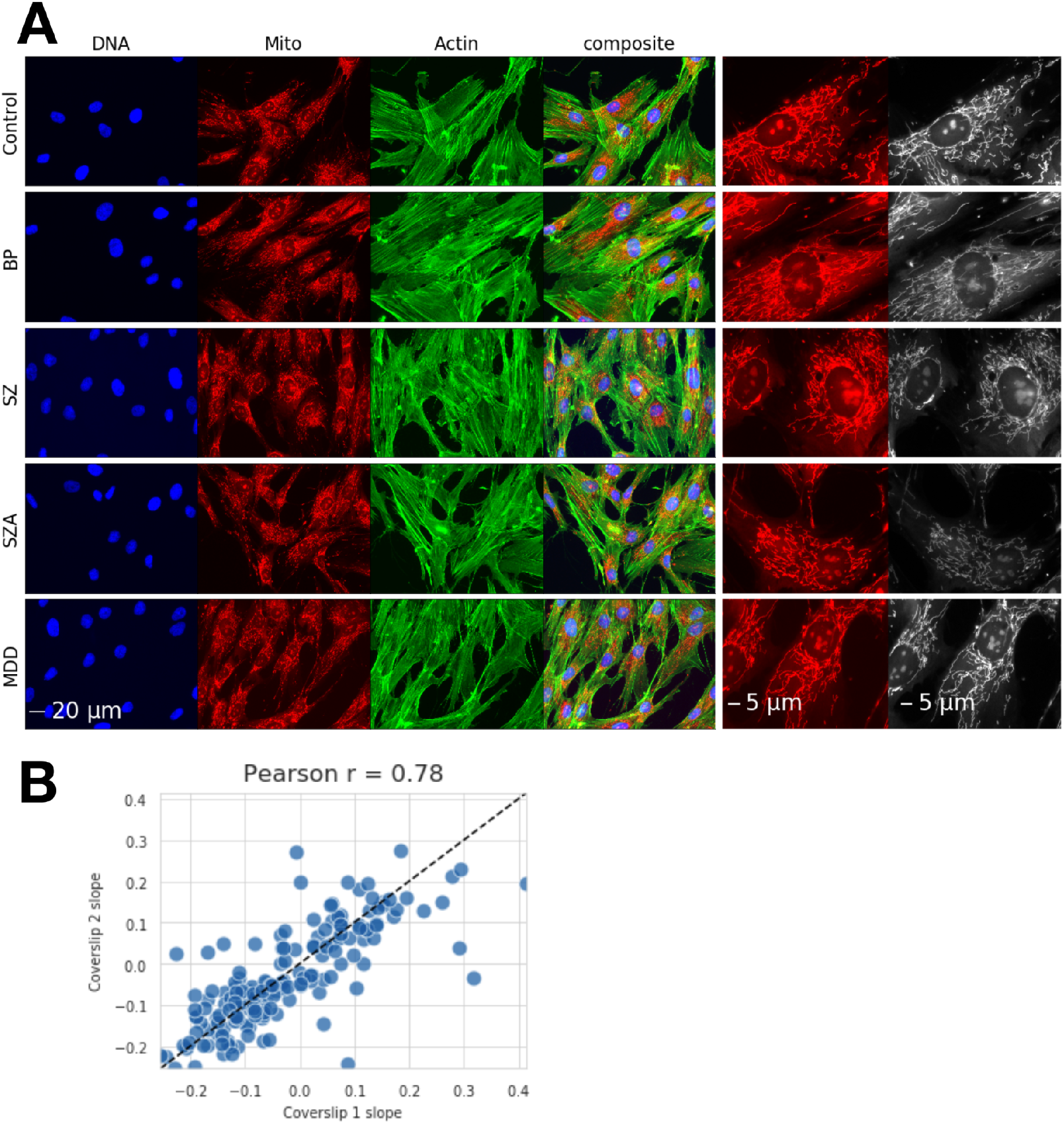
Image-based phenotypes associated with psychiatric conditions are subtle but reproducible. A) Fibroblasts that are stained for DNA, actin, and mitochondria, and labeled according to the patient category. Representative images were selected by identifying the subject closest to the median MITO-SLOPE for each group, then choosing the image from that subject with a MITO-SLOPE closest to the group median; the larger field of view shows the phenotype more clearly. Color images are clipped to their 0.5 and 99.5 percentile of their range and scaled to [0, 255] range just for visual clarity, given that the overall intensity of the staining is not the phenotype to be observed here. The grayscale images of mitochondria in the last column are scaled individually to the min/max, again just for display. Scale bar = 20 μm for the first four images in each row, which are full fields of view. Cropped images of MitoTracker in the last two columns have a scale bar of 5 μm. B) MITO-SLOPE values for the two replicate coverslips that were plated for each subject are shown, which yield a Pearson’s correlation coefficient (r) of 0.78.

**Supplementary Figure 3.**
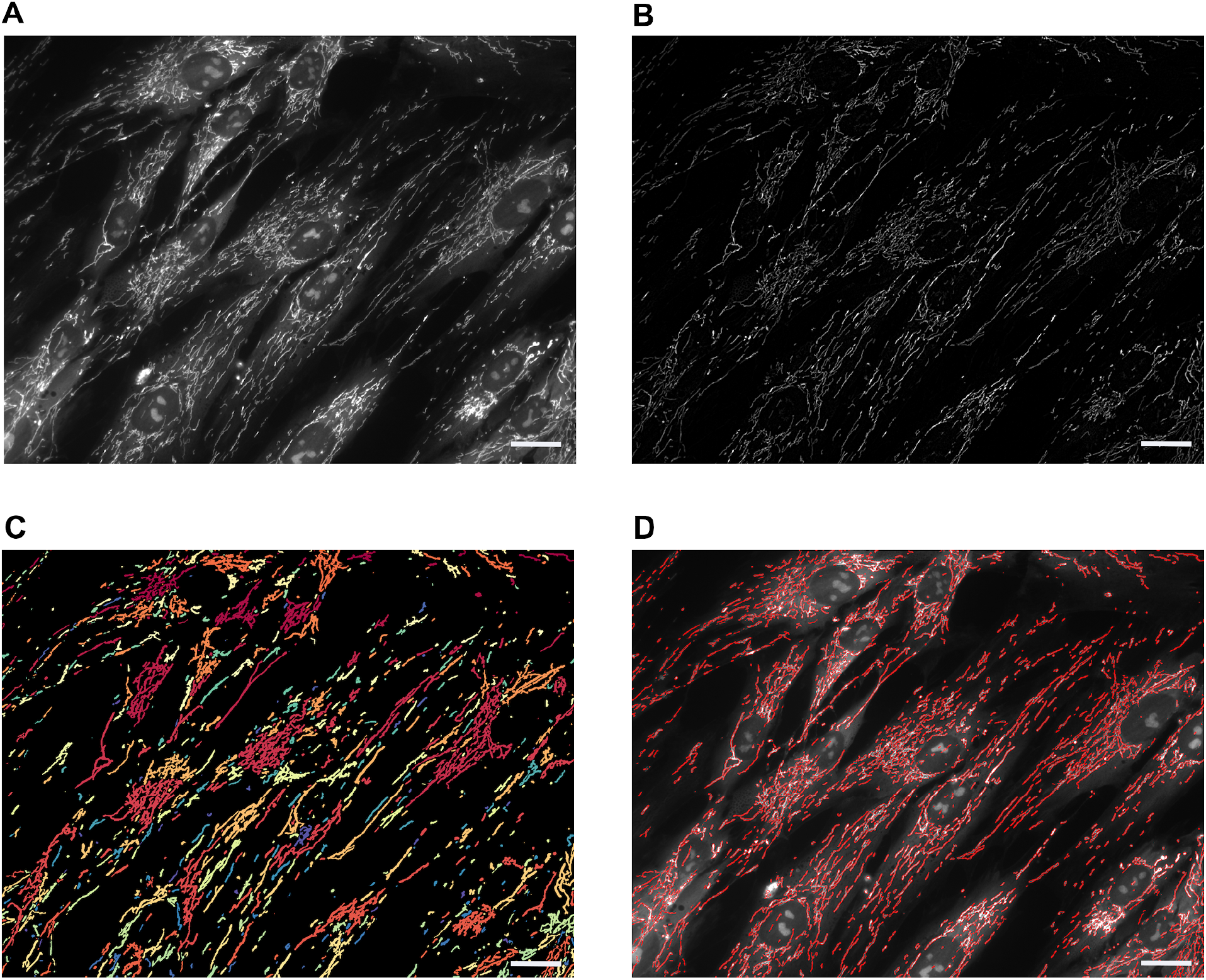
Mitochondrial segmentation is unsatisfactory for the study. Segmentation of mitochondria, even when optimized, do not appear to represent real, biologically meaningful boundaries for the cell type and imaging conditions of this study.

**Supplementary Figure 4.**
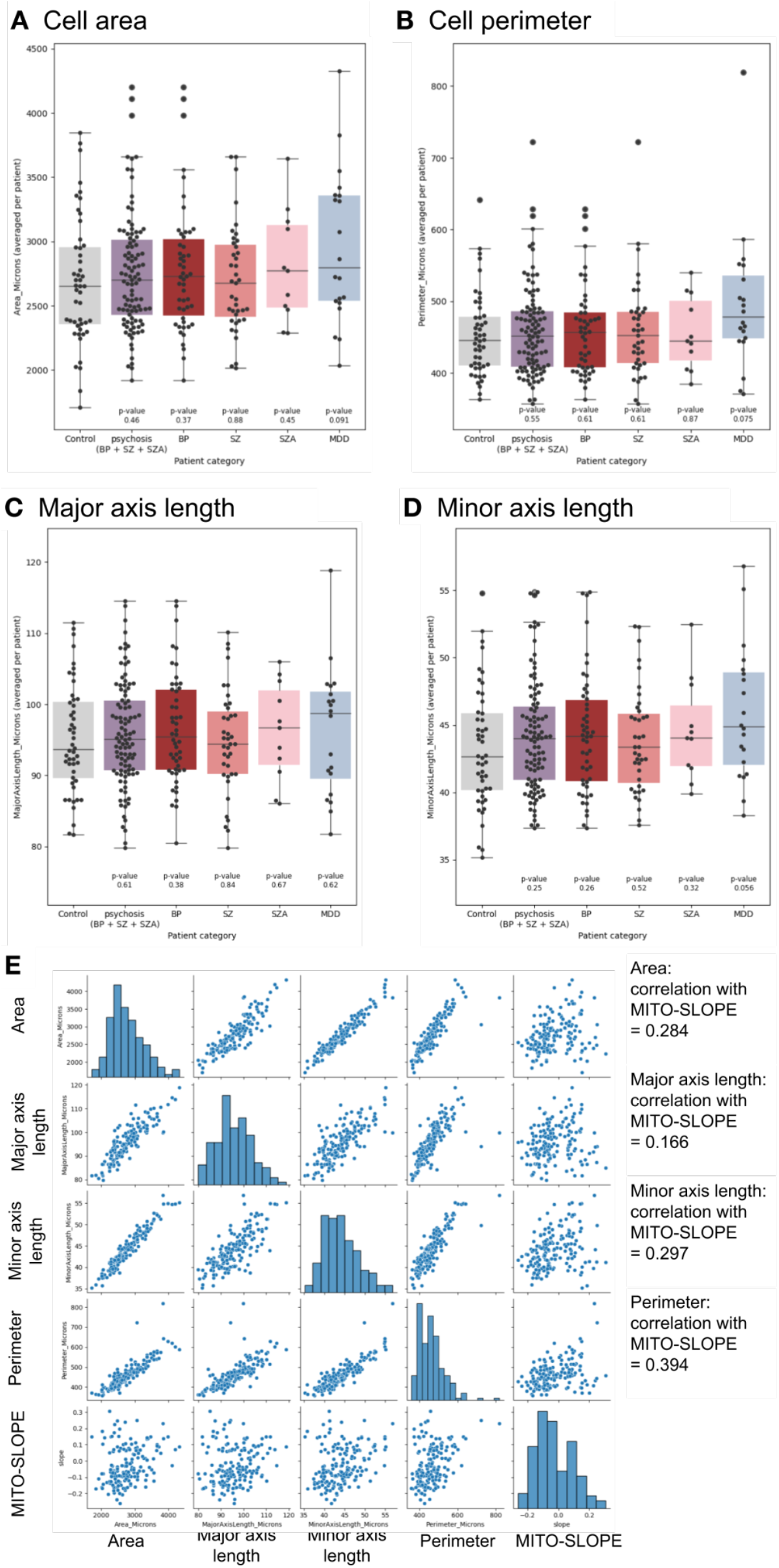
Various metrics of cell size do not distinguish various patient categories. (A. area, B. perimeter, C. major axis length, D. minor axis length). Data display is the same as in Figure 2D. E. Cell size metrics correlate strongly with each other but correlate less than 0.40 with MITO-SLOPE.

**Supplementary Figure 5.**
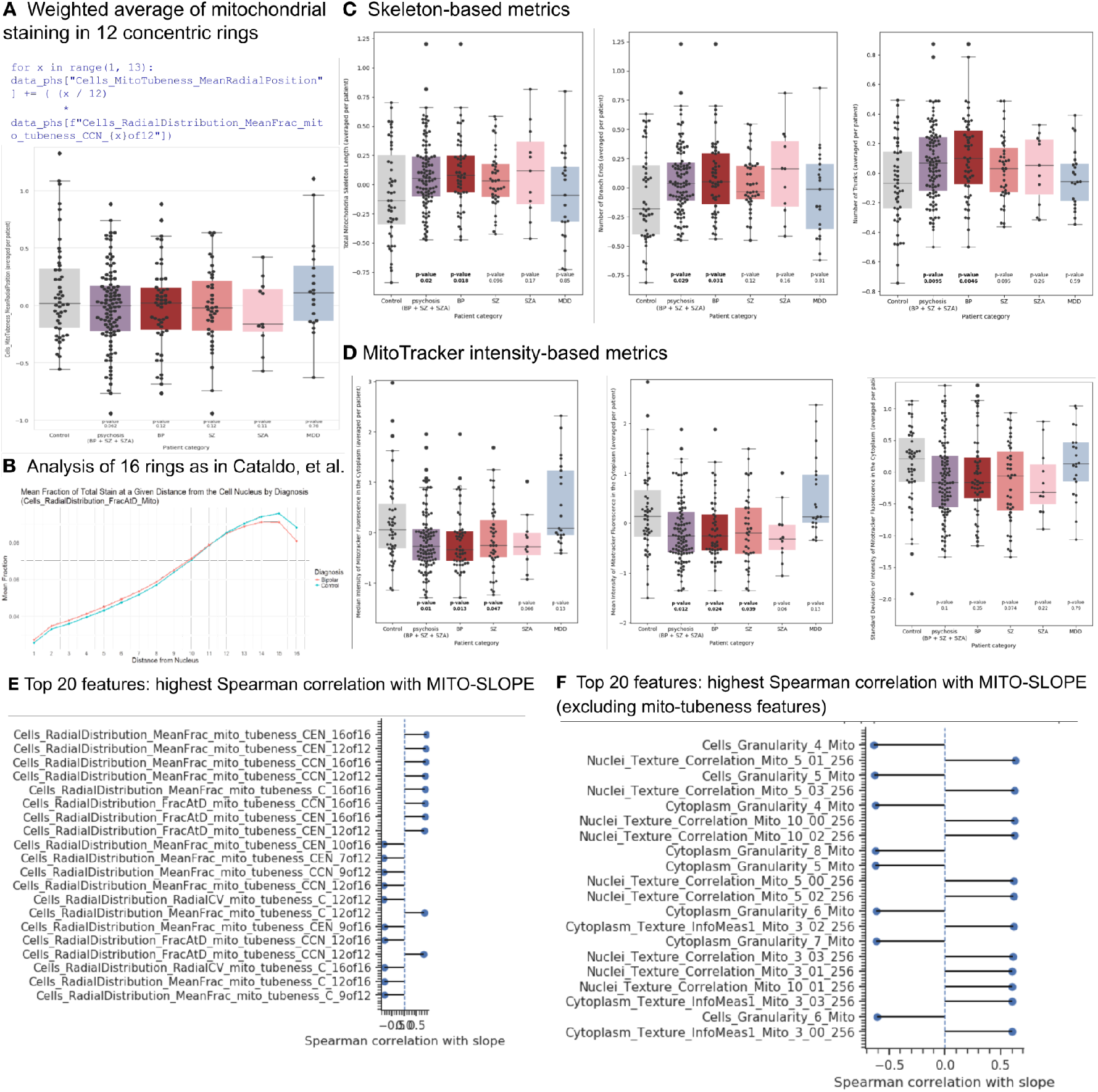
Various alternative metrics to characterize mitochondria. Plots for A, C, and D are as in Figure 2D except whiskers denote min/max values, except those values exceeding 1.5 x IQR are plotted as diamonds or larger circles. A) Weighted average of mitochondrial staining in 12 concentric rings following the formula shown. B) Analysis comparing the radial mitochondrial distribution across 16 rings between bipolar disorder and healthy control groups, as in Cataldo et al. (2010), using the feature Cells_RadialDistribution_FracAtD_Mito (the same feature we used for MITO-SLOPE except a larger number of bins). The area between mean profile curves was significantly greater than expected under a shared distribution (p=0.006), replicating prior results. C) Skeleton-based metrics involve thresholding and skeletonizing the mitochondrial network and using CellProfiler’s MeasureSkeleton module to measure various properties. This module measures the number of trunks and branches for each branching system in an image. The module takes a skeletonized image of the object plus previously identified seed objects (in this case, nuclei) and finds the number of trunks that emerge from the seed object and the number of branches along the trunks. Metrics shown are TotalObjectSkeletonLength (the length of all skeleton segments per object), NumberBranchEnds (the number of branch end-points, i.e, termini), and NumberTrunks (the number of trunks, i.e., branchpoints that lie within the seed objects). D) MitoTracker intensity-based metrics include the median, mean, and standard deviation of the intensity of MitoTracker staining in the cytoplasm, measured on a single-cell level. No segmentation of mitochondria was performed. E & F) Listing of features highly correlated with MITO-SLOPE.

**Supplementary Figure 6.**
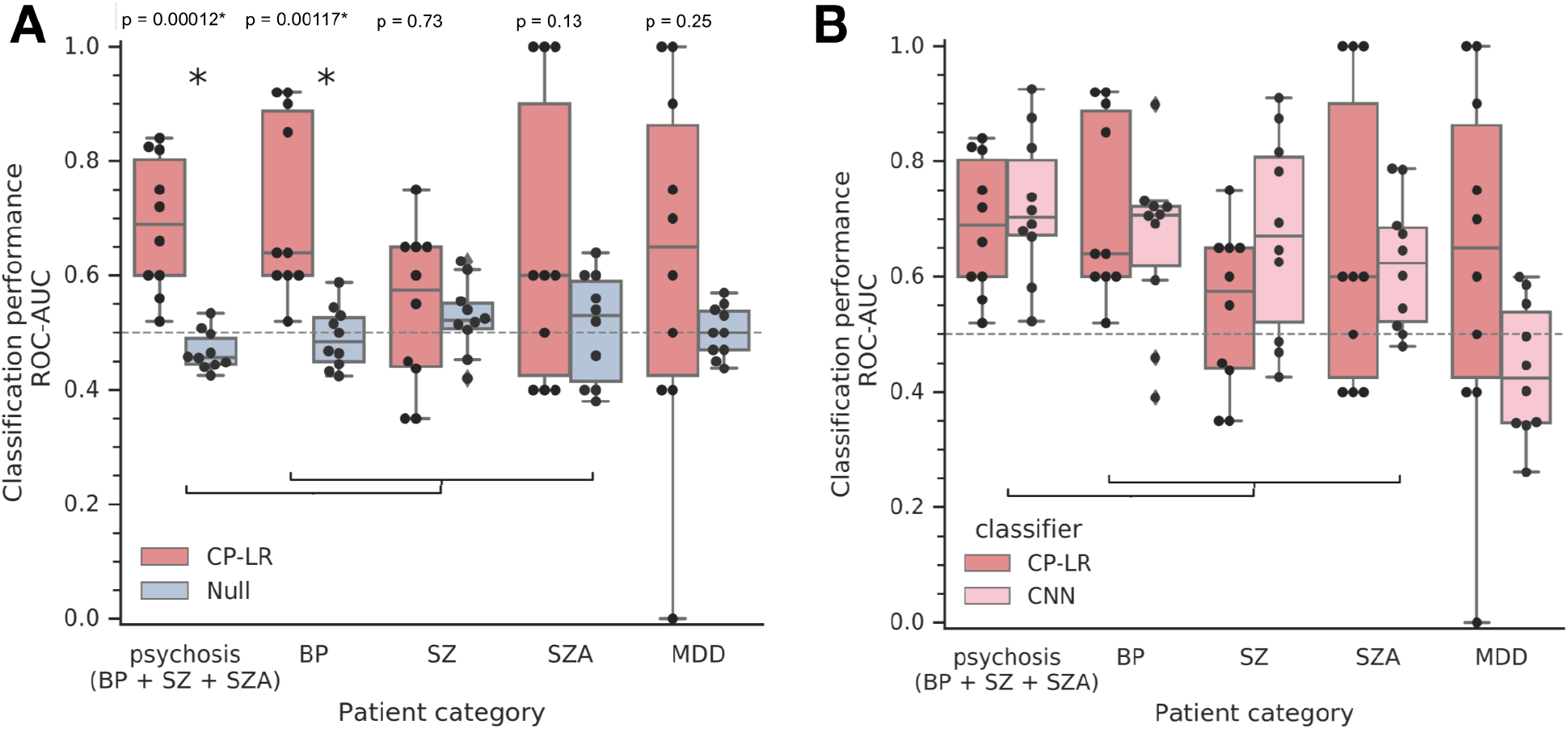
Machine learning / deep learning classification based on multiple mitochondrial features distinguishes patient group experiencing psychosis from healthy controls. A) Logistic Regression (LR) classifiers were trained on patient-level profiles constructed from mitochondrial features extracted using CellProfiler (CP) software (CP-LR). Patient categories include those with a primary diagnosis of bipolar disorder (BP), schizophrenia (SZ), schizoaffective disorder (SZA), and major depressive disorder (MDD). Psychosis encompasses patients in the BP, SZ, and SZA groups. Classifier performance is evaluated using 10-fold cross-validation, with the distribution of the area under the receiver operating characteristic curve (AUC-ROC) for each binary classification shown as individual data points on the box plots. The box plots represent the interquartile range (25th–75th percentile), with the median marked by a horizontal line; whiskers denote min/max values, except those values exceeding 1.5 x IQR are plotted as diamonds. The null distribution (blue) was generated by training classifiers on shuffled patient class labels. Most patient groups were not completely distinguishable from healthy controls in all folds of our 10-fold cross validation run, consistent with the overlap in patient groups seen with MITO-SLOPE. Statistically significant differences between observed and null classification performance (e.g., psychosis vs. healthy) were assessed using an independent two-sample t-test and are denoted by asterisks. B) Classification using CP-LR is shown as in A. For CNN, raw single-cell images of the mitochondrial channel were used for a deep learning classifier, a convolutional neural network (CNN). Patient categories include a primary diagnosis of bipolar disorder (BP), Schizophrenia (SZ), Schizoaffective disorder (SZA), and depression or major depressive disorder (MDD). Psychosis includes patients in classes BP, SZ, and SZA. Classifiers are evaluated by k-fold (k=10) cross validation and the distribution of area under the receiver operating characteristic curve (AUC-ROC) for each binary classification in each validation partition are reported as the data points comprising the box plots, which show 25th and 75th percentile, with the median marked as a line. Features used in the classification include all MitoTracker related features except those measuring correlation to other channels.

**Supplementary Figure 7.**
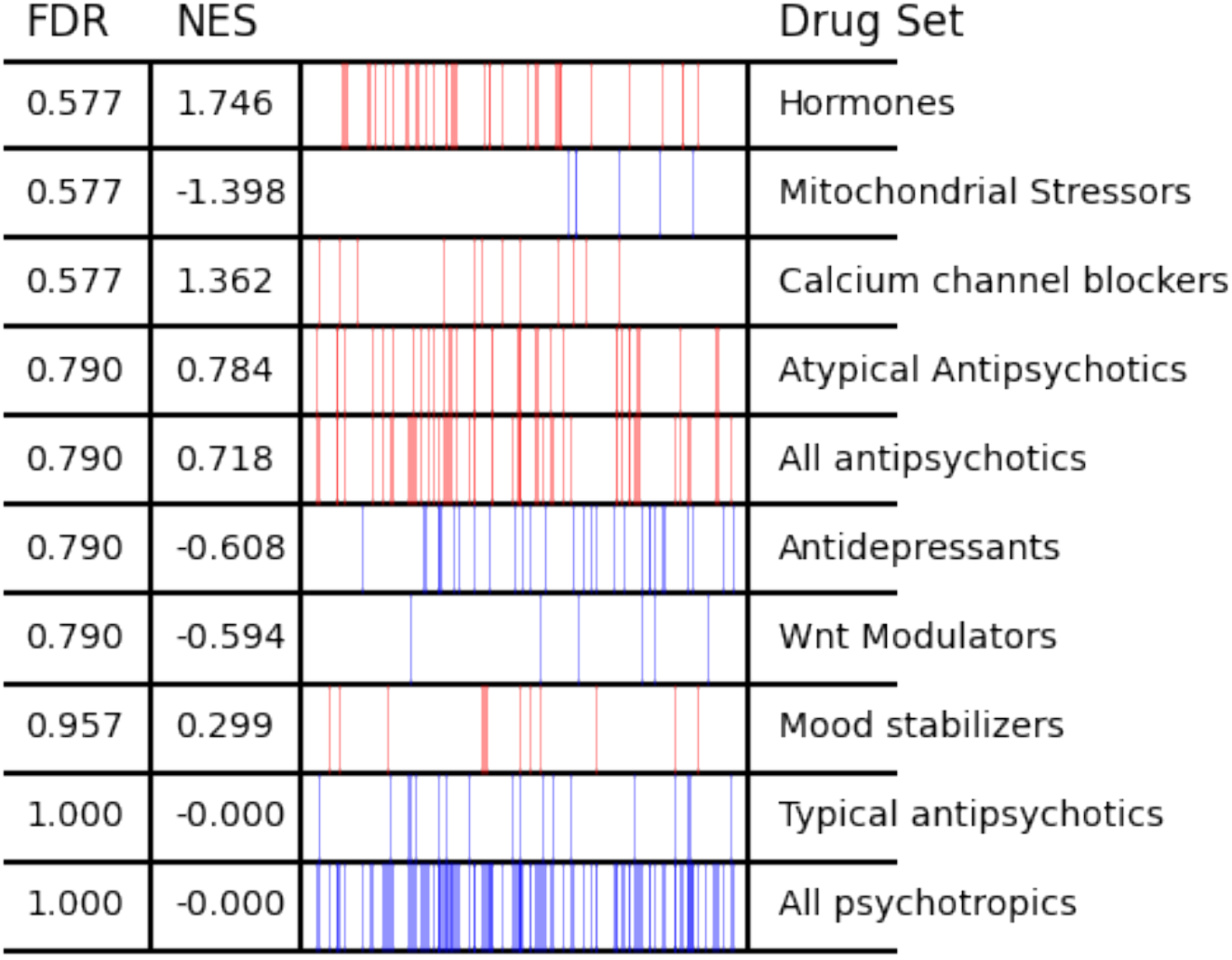
Existing classes of psychoactive drugs do not, overall, show evidence of impacting mitochondrial localization (all doses). Data is shown as in main Figure 3, but here with each compound-concentration shown as a separate data point and with all classes shown. As an example, although the data in the second row appears skewed to the right, it does not show statistically significant rank ordering in the assay because the Mitochondrial Stressors class contains only a single compound.

**Supplementary Figure 8.**
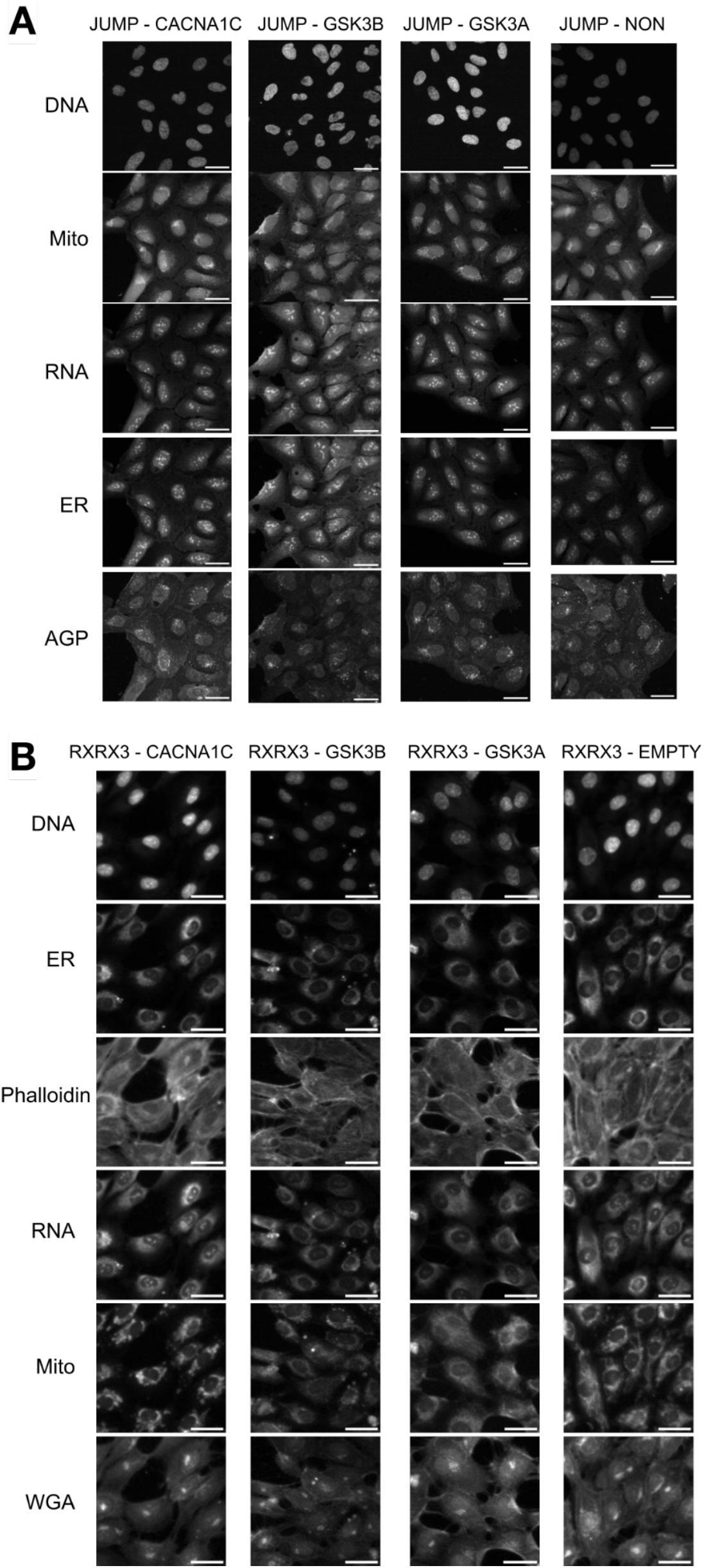
Mitochondrial localization impact for genes with known association with mitochondrial function or psychosis - all channels. An additional field of view is shown for Figure 5, with all channels shown as grayscale. AGP = actin, golgi, plasma membrane (as in Cell Painting); WGA = wheat germ agglutinin. Scale bar = 50 μm.

**Supplementary Figure 9.**
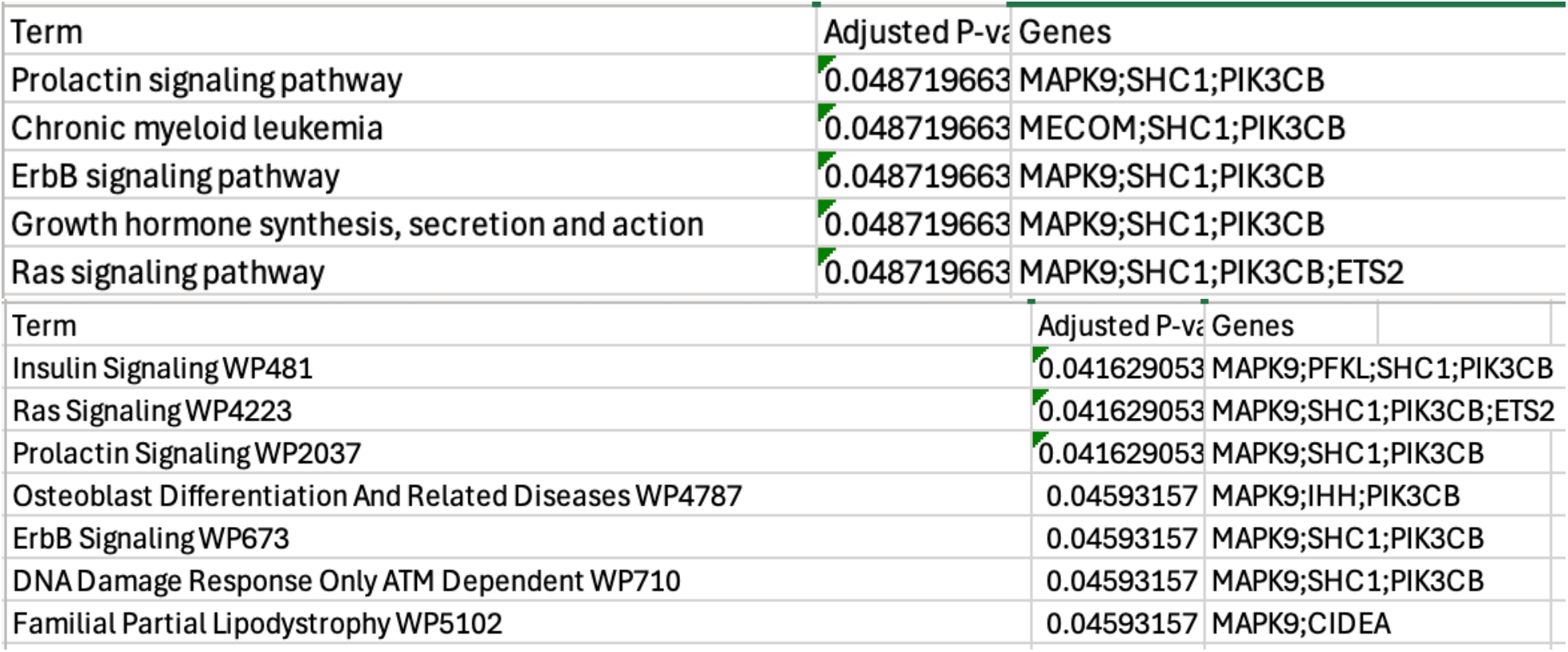
Enrichment analysis of the 39 overexpression hit genes using Enrichr. P-values are adjusted for multiple hypothesis testing across classes (but not across the 8 libraries/ontologies tested); the analysis used the tested genes as background. KEGG (top) and WikiPathways (bottom) are shown; the remaining six libraries/ontologies had no significant enrichments (Reactome, MSigDB, Hallmark sets, GO BP, GO CC, GO MF). Enrichment results files are available in the paper’s Github repository here: https://github.com/carpenter-singh-lab/2025_Haghighi_Mito/tree/main/results

**Supplementary Figure 10.**
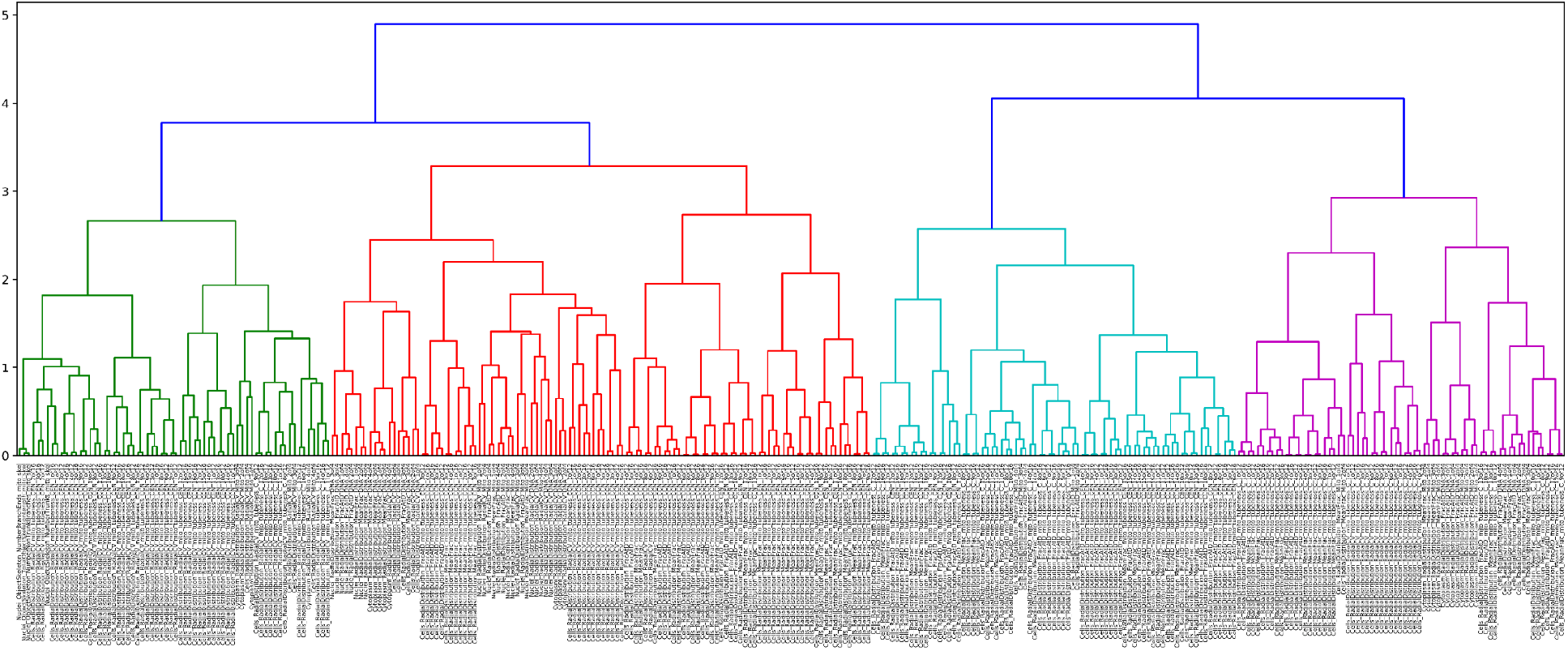
Hierarchical clustering dendrogram of mitochondrial channel features used for feature selection. The dendrogram illustrates clustering of 326 CellProfiler-extracted mitochondrial features based on Pearson correlation. A height threshold was selected to form the maximum number of feature clusters while ensuring that the resulting representative features had a Variance Inflation Factor (VIF) < 10, mitigating multicollinearity. One feature per cluster (the median feature) was selected to form a reduced, interpretable feature set of 45 representatives used for feature importance analysis in classification tasks. A high-resolution version of this figure is available here: https://github.com/carpenter-singh-lab/2025_Haghighi_Mito/blob/main/results/dendrogram_mito.pdf

**Supplementary Figure 11.**
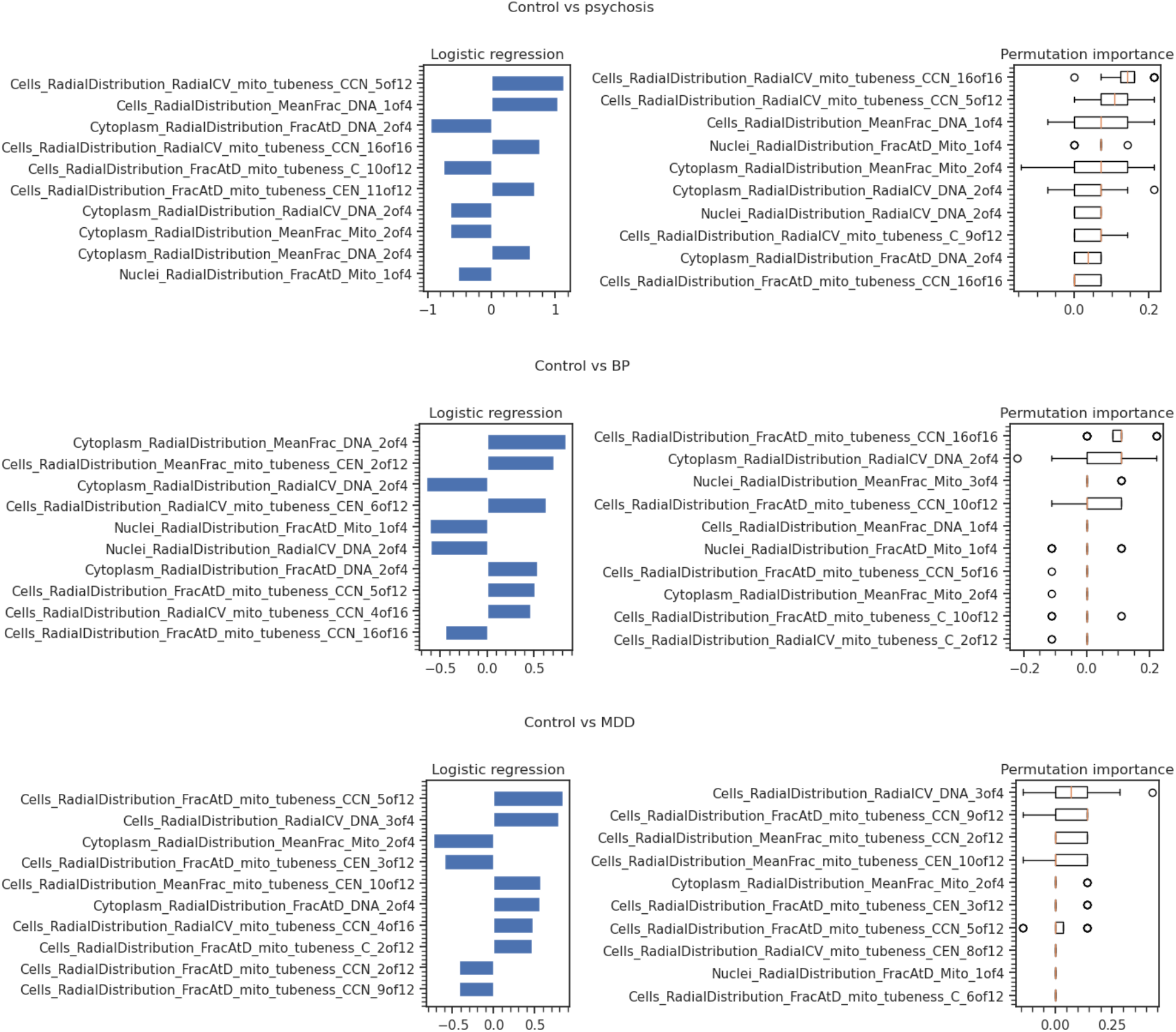
Feature importance analysis for distinguishing patient subgroups from controls using logistic regression models trained on representative CellProfiler mitochondrial features. For each binary classification task (Control vs. Psychosis, BP, SZ, SZA, or MDD), models were trained using a reduced set of 45 decorrelated features selected via hierarchical clustering to mitigate multicollinearity. Shown are the top 10 features ranked by absolute model coefficients (left) and permutation importance (right), for subgroups with AUC-ROC > 0.6. Results highlight texture and intensity-based mitochondrial features particularly those capturing radial intensity patterns as key contributors to classification performance.

## References

Al-Mehdi A-B, Pastukh VM, Swiger BM, et al (2012) Perinuclear mitochondrial clustering creates an oxidant-rich nuclear domain required for hypoxia-induced transcription. Sci Signal 5:ra47

Andreassen OA, Djurovic S, Thompson WK, et al (2013) Improved detection of common variants associated with schizophrenia by leveraging pleiotropy with cardiovascular-disease risk factors. Am J Hum Genet 92:197–209

Anon. (2021) Recursion and Bayer Expand Fibrosis Collaboration to Include Inferential Search Capabilities. In: Recursion website. https://ir.recursion.com/news-releases/news-release-details/recursion-and-bayer-expand-fibrosis-collaboration-include. Accessed 3 Mar 2025

Antony PMA, Kondratyeva O, Mommaerts K, et al (2020) Fibroblast mitochondria in idiopathic Parkinson’s disease display morphological changes and enhanced resistance to depolarization. Sci Rep 10:1569

Atkin TA, Brandon NJ, Kittler JT (2012) Disrupted in Schizophrenia 1 forms pathological aggresomes that disrupt its function in intracellular transport. Hum Mol Genet 21:2017–2028

Atkin TA, MacAskill AF, Brandon NJ, Kittler JT (2011) Disrupted in Schizophrenia-1 regulates intracellular trafficking of mitochondria in neurons. Mol Psychiatry 16:122– 4, 121

Benítez-King G, Valdés-Tovar M, Trueta C, et al (2016) The microtubular cytoskeleton of olfactory neurons derived from patients with schizophrenia or with bipolar disorder: Implications for biomarker characterization, neuronal physiology and pharmacological screening. Mol Cell Neurosci 73:84–95

Bessonova L, Ogden K, Doane MJ, et al (2020) The Economic Burden of Bipolar Disorder in the United States: A Systematic Literature Review. Clinicoecon Outcomes Res 12:481– 497

Blanchet L, Smeitink JAM, van Emst-de Vries SE, et al (2015) Quantifying small molecule phenotypic effects using mitochondrial morpho-functional fingerprinting and machine learning. Sci Rep 5:8035

Bray M-A, Gustafsdottir SM, Rohban MH, et al (2017) A dataset of images and morphological profiles of 30 000 small-molecule treatments using the Cell Painting assay. Gigascience 6:1–5

Brennand K, Savas JN, Kim Y, et al (2015) Phenotypic differences in hiPSC NPCs derived from patients with schizophrenia. Mol Psychiatry 20:361–368

Brown AS, Borgmann-Winter K, Hahn C-G, et al (2014) Increased stability of microtubules in cultured olfactory neuroepithelial cells from individuals with schizophrenia. Progress in Neuro-Psychopharmacology and Biological Psychiatry 48:252–258

Bulik-Sullivan B, Finucane HK, Anttila V, et al (2015) An atlas of genetic correlations across human diseases and traits. Nat Genet 47:1236–1241

Bulthuis EP, Adjobo-Hermans MJW, Willems PHGM, Koopman WJH (2019) Mitochondrial Morphofunction in Mammalian Cells. Antioxid Redox Signal 30:2066–2109

Carling PJ, Mortiboys H, Green C, et al (2020) Deep phenotyping of peripheral tissue facilitates mechanistic disease stratification in sporadic Parkinson’s disease. Prog Neurobiol 187:101772

Cataldo AM, McPhie DL, Lange NT, et al (2010) Abnormalities in mitochondrial structure in cells from patients with bipolar disorder. Am J Pathol 177:575–585

Celik S, Hütter J-C, Carlos SM, et al (2024) Building, benchmarking, and exploring perturbative maps of transcriptional and morphological data. PLoS Comput Biol 20:e1012463

Chan DC (2020) Mitochondrial Dynamics and Its Involvement in Disease. Annu Rev Pathol 15:235–259

Chandrasekaran SN, Ackerman J, Alix E, et al (2023) JUMP Cell Painting dataset: morphological impact of 136,000 chemical and genetic perturbations. bioRxiv 2023.03.23.534023

Chandrasekaran SN, Alix E, Arevalo J, et al (2024) Morphological map of under- and over-expression of genes in human cells. bioRxiv

Chandrasekaran SN, Ceulemans H, Boyd JD, Carpenter AE (2020) Image-based profiling for drug discovery: due for a machine-learning upgrade? Nat Rev Drug Discov 1–15

Chen H, Lu M, Lyu Q, et al (2024) Mitochondrial dynamics dysfunction: Unraveling the hidden link to depression. Biomed Pharmacother 175:116656

Chu C-H, Tseng W-W, Hsu C-M, Wei A-C (2022) Image analysis of the mitochondrial network morphology with applications in cancer research. Front Phys 10:. 10.3389/fphy.2022.855775

Cohen BM, Öngür D (2018) The urgent need for more research on bipolar depression. Lancet Psychiatry 5:e29–e30

Corena-McLeod M, Walss-Bass C, Oliveros A, et al (2013) New model of action for mood stabilizers: phosphoproteome from rat pre-frontal cortex synaptoneurosomal preparations. PLoS One 8:e52147

Cuccarese MF, Earnshaw BA, Heiser K, et al (2020) Functional immune mapping with deep-learning enabled phenomics applied to immunomodulatory and COVID-19 drug discovery. bioRxiv 2020.08.02.233064

Culmsee C, Michels S, Scheu S, et al (2018) Mitochondria, Microglia, and the Immune System-How Are They Linked in Affective Disorders? Front Psychiatry 9:739

Cuperfain AB, Zhang ZL, Kennedy JL, Gonçalves VF (2018) The Complex Interaction of Mitochondrial Genetics and Mitochondrial Pathways in Psychiatric Disease. Mol Neuropsychiatry 4:52–69

Ding JB, Hu K (2021) Cigarette smoking and schizophrenia: Etiology, clinical, pharmacological, and treatment implications. Schizophr Res Treatment 2021:7698030

Erhardt S, Schwieler L, Imbeault S, Engberg G (2017) The kynurenine pathway in schizophrenia and bipolar disorder. Neuropharmacology 112:297–306

Fay MM, Kraus O, Victors M, et al (2023) RxRx3: Phenomics Map of Biology. bioRxiv 2023.02.07.527350

Fischer NC, Friedman V, Martinez-Reyes MA, et al (2022) The ANC-1 (Nesprin-1/2) organelle-anchoring protein functions through mitochondria to polarize axon growth in response to SLT-1. PLoS Genet 18:e1010521

Flippo KH, Strack S (2017) An emerging role for mitochondrial dynamics in schizophrenia. Schizophr Res 187:26–32

Giménez-Palomo A, Dodd S, Anmella G, et al (2021) The Role of Mitochondria in Mood Disorders: From Physiology to Pathophysiology and to Treatment. Front Psychiatry 12:546801

Goering R, Arora A, Pockalny MC, Taliaferro JM (2023) RNA localization mechanisms transcend cell morphology. Elife 12:e80040

Gray JA, Roth BL (2007) The pipeline and future of drug development in schizophrenia. Mol Psychiatry 12:904–922

Grotzinger AD, Singh K, Miller-Fleming TW, et al (2023) Transcriptome-Wide Structural Equation Modeling of 13 Major Psychiatric Disorders for Cross-Disorder Risk and Drug Repurposing. JAMA Psychiatry. 10.1001/jamapsychiatry.2023.1808

Harrison PJ, Colbourne L, Harrison CH (2020) The neuropathology of bipolar disorder: systematic review and meta-analysis. Mol Psychiatry 25:1787–1808

Harwig MC, Viana MP, Egner JM, et al (2018) Methods for imaging mammalian mitochondrial morphology: A prospective on MitoGraph. Anal Biochem 552:81–99

Hebb ALO, Robertson HA (2007) Role of phosphodiesterases in neurological and psychiatric disease. Curr Opin Pharmacol 7:86–92

Heffner JL, Strawn JR, DelBello MP, et al (2011) The co-occurrence of cigarette smoking and bipolar disorder: phenomenology and treatment considerations: Cigarette smoking and bipolar disorder. Bipolar Disord 13:439–453

Hemel IMGM, Engelen BPH, Luber N, Gerards M (2021) A hitchhiker’s guide to mitochondrial quantification. Mitochondrion 59:216–224

Hirabayashi Y, Lewis TL, Du Y, et al (2024) Most axonal mitochondria in cortical pyramidal neurons lack mitochondrial DNA and consume ATP. bioRxiv. 10.1101/2024.02.12.579972

Hung CL-K, Maiuri T, Bowie LE, et al (2018) A patient-derived cellular model for Huntington’s disease reveals phenotypes at clinically relevant CAG lengths. Mol Biol Cell 29:2809–2820

Jenkins BC, Neikirk K, Katti P, et al (2024) Mitochondria in disease: changes in shapes and dynamics. Trends Biochem Sci 49:346–360

Jiang M, Yan W, Zhang Y, et al (2023) Phosphodiesterase and psychiatric disorders: a two-sample Mendelian randomization study. J Transl Med 21:560

Jope RS, Roh M-S (2006) Glycogen synthase kinase-3 (GSK3) in psychiatric diseases and therapeutic interventions. Curr Drug Targets 7:1421–1434

Kadakia A, Catillon M, Fan Q, et al (2022) The economic burden of schizophrenia in the United States. J Clin Psychiatry 83:. 10.4088/JCP.22m14458

Kessi M, Chen B, Pan L, et al (2023) Disruption of mitochondrial and lysosomal functions by human CACNA1C variants expressed in HEK 293 and CHO cells. Front Mol Neurosci 16:1209760

Kuperberg M, Greenebaum SLA, Nierenberg AA (2021) Targeting Mitochondrial Dysfunction for Bipolar Disorder. In: Young AH, Juruena MF (eds) Bipolar Disorder: From Neuroscience to Treatment. Springer International Publishing, Cham, pp 61–99

Kwon S-K, Sando R 3rd, Lewis TL, et al (2016) LKB1 regulates mitochondria-dependent presynaptic calcium clearance and neurotransmitter release properties at excitatory synapses along cortical axons. PLoS Biol 14:e1002516

Lachmann A, Xie Z, Ma’ayan A (2022) blitzGSEA: efficient computation of gene set enrichment analysis through gamma distribution approximation. Bioinformatics 38:2356–2357

Lamb J, Crawford ED, Peck D, et al (2006) The Connectivity Map: using gene-expression signatures to connect small molecules, genes, and disease. Science 313:1929–1935

Leonard AP, Cameron RB, Speiser JL, et al (2015) Quantitative analysis of mitochondrial morphology and membrane potential in living cells using high-content imaging, machine learning, and morphological binning. Biochim Biophys Acta 1853:348–360

Lin C, Zhang X, Jin H (2022) The Societal Cost of Schizophrenia: An Updated Systematic Review of Cost-of-Illness Studies. Pharmacoeconomics. 10.1007/s40273-022-01217-8

Lones MA (2021) How to avoid machine learning pitfalls: a guide for academic researchers. arXiv [cs.LG]

Mandal A, Drerup CM (2019) Axonal Transport and Mitochondrial Function in Neurons. Front Cell Neurosci 13:373

Marchisella F, Coffey ET, Hollos P (2016) Microtubule and microtubule associated protein anomalies in psychiatric disease. Cytoskeleton (Hoboken) 73:596–611

Marques AP, Resende R, Silva DF, et al (2021) Mitochondrial Alterations in Fibroblasts of Early Stage Bipolar Disorder Patients. Biomedicines 9:. 10.3390/biomedicines9050522

McDiarmid AH, Gospodinova KO, Elliott RJR, et al (2023) Morphological profiling by Cell Painting in human neural progenitor cells classifies hit compounds in a pilot drug screen for Alzheimer’s disease. bioRxiv 2023.01.16.523559

McIntyre RS, Berk M, Brietzke E, et al (2020) Bipolar disorders. Lancet 396:1841–1856

McPhie DL, Nehme R, Ravichandran C, et al (2018) Oligodendrocyte differentiation of induced pluripotent stem cells derived from subjects with schizophrenias implicate abnormalities in development. Transl Psychiatry 8:230

McPhie ML, Bridgman AC, Kirchhof MG (2021) A Review of Skin Disease in Schizophrenia. Dermatology 237:248–261

McQuin C, Goodman A, Chernyshev V, et al (2018) CellProfiler 3.0: Next-generation image processing for biology. PLoS Biol 16:e2005970

Mertens J, Wang Q-W, Kim Y, et al (2015) Differential responses to lithium in hyperexcitable neurons from patients with bipolar disorder. Nature 527:95–99

Michels S, Ganjam GK, Martins H, et al (2018) Downregulation of the psychiatric susceptibility gene Cacna1c promotes mitochondrial resilience to oxidative stress in neuronal cells. Cell Death Discov 4:54

Moon AL, Haan N, Wilkinson LS, et al (2018) CACNA1C: Association with psychiatric disorders, behavior, and neurogenesis. Schizophr Bull 44:958–965

Mortiboys H, Johansen KK, Aasly JO, Bandmann O (2010) Mitochondrial impairment in patients with Parkinson disease with the G2019S mutation in LRRK2. Neurology 75:2017–2020

Mortiboys H, Thomas KJ, Koopman WJH, et al (2008) Mitochondrial function and morphology are impaired in parkin-mutant fibroblasts. Ann Neurol 64:555–565

Munakata K, Tanaka M, Mori K, et al (2004) Mitochondrial DNA 3644T-->C mutation associated with bipolar disorder. Genomics 84:1041–1050

Murphy LC, Millar JK (2017) Regulation of mitochondrial dynamics by DISC1, a putative risk factor for major mental illness. Schizophr Res 187:55–61

Neumann C, Shankaran H, Nadella K, et al (2024) Abstract 7133: Phenomics-enabled discovery and optimization of small-molecule RBM39 degraders as an alternative to CDK12 targeting in high-grade serous ovarian cancer (HGSOC). Cancer Res 84:7133– 7133

Ni P, Ma Y, Chung S (2024) Mitochondrial dysfunction in psychiatric disorders. Schizophr Res 273:62–77

Olesen MA, Villavicencio-Tejo F, Quintanilla RA (2022) The use of fibroblasts as a valuable strategy for studying mitochondrial impairment in neurological disorders. Transl Neurodegener 11:36

Panchision DM (2016) Concise Review: Progress and Challenges in Using Human Stem Cells for Biological and Therapeutics Discovery: Neuropsychiatric Disorders. Stem Cells 34:523–536

Papageorgiou MP, Filiou MD (2024) Mitochondrial dynamics and psychiatric disorders: The missing link. Neurosci Biobehav Rev 165:105837

Payne T, Burgess T, Bradley S, et al (2024) Multimodal assessment of mitochondrial function in Parkinson’s disease. Brain 147:267–280

Pereira C, Chavarria V, Vian J, et al (2018) Mitochondrial Agents for Bipolar Disorder. Int J Neuropsychopharmacol 21:550–569

Perrottelli A, Flavio Marzocchi F, Caporusso E, et al Advances in the understanding of the pathophysiology of schizophrenia and bipolar disorder through induced pluripotent stem cell models

Plitman E, Iwata Y, Caravaggio F, et al (2017) Kynurenic acid in schizophrenia: A systematic review and meta-analysis. Schizophr Bull 43:764–777

Qi HAO, Qiu T, Chu PK, et al (2018) Nuclear staining with MitoTracker® Orange CMTMRos (Thermo Fisher Scientific, cat. no. M7510)? In: ResearchGate. https://www.researchgate.net/post/Nuclear-staining-with-MitoTrackerR-Orange-CMTMRos-Thermo-Fisher-Scientific-cat-no-M7510. Accessed 10 Feb 2026

Qian X, Song H, Ming G-L (2019) Brain organoids: advances, applications and challenges. Development 146:. 10.1242/dev.166074

Ripoll N, Bronnec M, Bourin M (2004) Nicotinic receptors and schizophrenia. Curr Med Res Opin 20:1057–1074

Roberts RC (2021) Mitochondrial dysfunction in schizophrenia: With a focus on postmortem studies. Mitochondrion 56:91–101

Robicsek O, Karry R, Petit I, et al (2013) Abnormal neuronal differentiation and mitochondrial dysfunction in hair follicle-derived induced pluripotent stem cells of schizophrenia patients. Mol Psychiatry 18:1067–1076

Rohban MH, Fuller AM, Tan C, et al (2022) Virtual screening for small-molecule pathway regulators by image-profile matching. Cell Syst 13:724–736.e9

Rosenfeld M, Brenner-Lavie H, Ari SG-B, et al (2011) Perturbation in mitochondrial network dynamics and in complex I dependent cellular respiration in schizophrenia. Biol Psychiatry 69:980–988

Sadeghi MA, Nassireslami E, Yousefi Zoshk M, et al (2023) Phosphodiesterase inhibitors in psychiatric disorders. Psychopharmacology (Berl) 240:1201–1219

Sato Y, Nakajima S, Shiraga N, et al (1998) Three-dimensional multi-scale line filter for segmentation and visualization of curvilinear structures in medical images. Med Image Anal 2:143–168

Scaini G, Andrews T, Lima CNC, et al (2021) Mitochondrial dysfunction as a critical event in the pathophysiology of bipolar disorder. Mitochondrion 57:23–36

Scaini G, Fries GR, Valvassori SS, et al (2017) Perturbations in the apoptotic pathway and mitochondrial network dynamics in peripheral blood mononuclear cells from bipolar disorder patients. Transl Psychiatry 7:e1111

Scaini G, Rezin GT, Carvalho AF, et al (2016) Mitochondrial dysfunction in bipolar disorder: Evidence, pathophysiology and translational implications. Neurosci Biobehav Rev 68:694–713

Schiff L, Migliori B, Chen Y, et al (2022) Integrating deep learning and unbiased automated high-content screening to identify complex disease signatures in human fibroblasts. Nat Commun 13:1590

Sklar P, Smoller JW, Fan J, et al (2008) Whole-genome association study of bipolar disorder. Mol Psychiatry 13:558–569

Smith GA, Jansson J, Rocha EM, et al (2016) Fibroblast biomarkers of sporadic Parkinson’s disease and LRRK2 kinase inhibition. Mol Neurobiol 53:5161–5177

Snelleksz M, Dean B (2021) Lower levels of tubulin alpha 1b in the frontal pole in schizophrenia supports a role for changed cytoskeletal dynamics in the aetiology of the disorder. Psychiatry Res 303:114096

Snitow ME, Bhansali RS, Klein PS (2021) Lithium and therapeutic targeting of GSK-3. Cells 10:255

Stępnicki P, Kondej M, Kaczor AA (2018) Current Concepts and Treatments of Schizophrenia. Molecules 23:. 10.3390/molecules23082087

Stirling DR, Swain-Bowden MJ, Lucas AM, et al (2021) CellProfiler 4: improvements in speed, utility and usability. BMC Bioinformatics 22:433

Subramanian A, Tamayo P, Mootha VK, et al (2005) Gene set enrichment analysis: a knowledge-based approach for interpreting genome-wide expression profiles. Proc Natl Acad Sci U S A 102:15545–15550

Suh BK, Lee S-A, Park C, et al (2021) Schizophrenia-associated dysbindin modulates axonal mitochondrial movement in cooperation with p150glued. Mol Brain 14:14

Sutton LP, Rushlow WJ (2011) The effects of neuropsychiatric drugs on glycogen synthase kinase-3 signaling. Neuroscience 199:116–124

Tan H-Y, Cho H, Lee LP (2021) Human mini-brain models. Nat Biomed Eng 5:11–25

Tegtmeyer M, Liyanage D, Han Y, et al (2024) Combining NeuroPainting with transcriptomics reveals cell-type-specific morphological and molecular signatures of the 22q11.2 deletion. bioRxivorg

Teves JMY, Bhargava V, Kirwan KR, et al (2017) Parkinson’s disease skin fibroblasts display signature alterations in growth, redox homeostasis, mitochondrial function, and autophagy. Front Neurosci 11:737

Theeuwes WF, Gosker HR, Langen RCJ, et al (2018) Inactivation of glycogen synthase kinase 3β (GSK-3β) enhances mitochondrial biogenesis during myogenesis. Biochim Biophys Acta Mol Basis Dis 1864:2913–2926

Thomas RA, Cai E, Reintsch W, et al (2023) A deep learning convolutional neural network distinguishes neuronal models of Parkinson’s disease from matched controls. bioRxiv 2023.11.23.568499

Thomsen MS, Weyn A, Mikkelsen JD (2011) Hippocampal α7 nicotinic acetylcholine receptor levels in patients with schizophrenia, bipolar disorder, or major depressive disorder: α7 nAChR in mental disorders. Bipolar Disord 13:701–707

Trubetskoy V, Pardiñas AF, Qi T, et al (2022) Mapping genomic loci implicates genes and synaptic biology in schizophrenia. Nature 604:502–508

Vaccaro V, Devine MJ, Higgs NF, Kittler JT (2017) Miro1-dependent mitochondrial positioning drives the rescaling of presynaptic Ca2+ signals during homeostatic plasticity. EMBO Rep 18:231–240

Varidaki A, Hong Y, Coffey ET (2018) Repositioning microtubule stabilizing drugs for brain disorders. Front Cell Neurosci 12:226

Wali G, Berkovsky S, Whiten DR, et al (2021) Single cell morphology distinguishes genotype and drug effect in Hereditary Spastic Paraplegia. Sci Rep 11:16635

Wali G, Li Y, Liyanage E, et al (2023) Pharmacological rescue of mitochondrial and neuronal defects in SPG7 hereditary spastic paraplegia patient neurons using high throughput assays. Front Neurosci 17:1231584

Wawer MJ, Li K, Gustafsdottir SM, et al (2014) Toward performance-diverse small-molecule libraries for cell-based phenotypic screening using multiplexed high-dimensional profiling. Proc Natl Acad Sci U S A 111:10911–10916

Way GP, Natoli T, Adeboye A, et al (2022) Morphology and gene expression profiling provide complementary information for mapping cell state. Cell Syst 13:911–923.e9

Whelan DR, Bell TDM (2015) Image artifacts in single molecule localization microscopy: why optimization of sample preparation protocols matters. Sci Rep 5:7924

Wu Y, Chen M, Jiang J (2019) Mitochondrial dysfunction in neurodegenerative diseases and drug targets via apoptotic signaling. Mitochondrion 49:35–45

Yang K, Chen Z, Gao J, et al (2017) The key roles of GSK-3β in regulating mitochondrial activity. Cell Physiol Biochem 44:1445–1459

Yang SJ, Lipnick SL, Makhortova NR, et al (2019) Applying Deep Neural Network Analysis to High-Content Image-Based Assays. SLAS Discovery 24:829–841

Yatchenko Y, Ben-Shachar D (2021) Update of mitochondrial network analysis by imaging: Proof of technique in schizophrenia. Methods Mol Biol 2277:187–201

Zanfardino P, Doccini S, Santorelli FM, Petruzzella V (2021) Tackling Dysfunction of Mitochondrial Bioenergetics in the Brain. Int J Mol Sci 22:. 10.3390/ijms22158325

