## Supplementary Table 1 for "Identifying and targeting abnormal mitochondrial localization associated with psychosis"

**Table . Subject Demographics, Comorbid Conditions and Medications at time of Biopsy**

| **Dx** | **Sex** | **Age** | **Race** | **Substance use** | **Comorbid Psych Dx** | **Comorbid Med Dx** | **Medications (as reported)** | **Medications (generic name)** |
| --- | --- | --- | --- | --- | --- | --- | --- | --- |
| Control | M | 55 | White | None | None | Arthritis | None | None |
| Control | M | 37 | White | None | None | None | None | None |
| Control | M | 24 | White | None | None | None | None | None |
| Control | M | 23 | White | None | None | None | None | None |
| Control | M | 24 | White | None | None | None | None | None |
| Control | F | 65 | White | None | None | None | None | None |
| Control | F | 27 | Asian | None | None | None | None | None |
| Control | M | 31 | White | None | None | None | None | None |
| Control | F | 44 | White | None | None | Urolithiasis (Urostomy), Cholecystectomy | Necon | ethinyl estradiol/norethindrone |
| Control | F | 52 | White | None | None | None | Vagifem cream, multivit, calcium, fish oil | estradiol vaginal cream, multivitamin, calcium, fish oil |
| Control | M | 26 | White/  Asian | None | None | None | None | None |
| Control | F | 54 | White | None | None | None | vit D, calcium, B12 | vitamin D, calcium, vitamin B12 |
| Control | F | 31 | White | None | None | None | no meds | no meds |
| Control | M | 58 | White | None | None | Stage 3 Melanoma 1 yr ago, Acute Pancreatitis,  Cholecystectomy 6 yrs ago | vit D | vitamin D |
| Control | M | 49 | White | None | None | None | None | None |
| Control | M | 41 | White | None | None | None | Tylenol prn | acetaminophen prn |
| Control | M | 40 | Black | None | None | None | None | None |
| Control | M | 42 | White | None | None | None | None | None |
| Control | F | 39 | White | None | None | None | None | None |
| Control | M | 45 | White | None | None | None | Zocor 40 mg/day | simvastatin 40 mg/day |
| Control | F | 24 | White | None | None | Pituitary Adenoma (no Sx) | multivit | multivitamin |
| Control | M | 24 | White | None | None | None | multivit | multivitamin |
| Control | M | 20 | Black | None | None | None | None | None |
| Control | M | 25 | White | None | None | None | None | None |
| Control | M | 21 | Asian | None | None | None | None | None |
| Control | M | 43 | White | None | None | None | None | None |
| Control | M | 23 | White | None | None | None | None | None |
| Control | M | 22 | White | None | None | None | None | None |
| Control | F | 32 | White | None | None | None | None | None |
| Control | F | 36 | Black | None | None | None | fish oil, multivit | fish oil, multivitamin |
| Control | F | 36 | White | None | None | None | Birth control pill | oral contraceptive pill |
| Control | M | 42 | Asian | None | None | None | None | None |
| Control | F | 22 | White | None | None | None | None | None |
| Control | M | 23 | White | None | None | None | None | None |
| Control | M | 57 | White | None | None | None | Lipitor 20, aspirin 81, multivit | atorvastatin 20, aspirin 81, multivitamin |
| Control | M | 31 | Black | None | None | None | None | None |
| Control | F | 48 | White | None | None | Hypertension | Cardizem, Yaz, multivit | diltiazem, drospirenone/ethinyl estradiol, multivitamin |
| Control | M | 52 | White | None | None | None | aspirin, multivit | aspirin, multivitamin |
| Control | M | 35 | White | None | None | None | None | None |
| Control | M | 21 | White | None | None | None | None | None |
| Control | M | 21 | SE Asian | None | None | None | None | None |
| Control | F | 52 | White | None | None | None | None | None |
| Control | F | 54 | White | None | None | None | calcium, vit D | calcium, vitamin D |
| Control | F | 31 | Black | None | None | None | None | None |
| Control | M | 24 | White | None | None | None | None | None |
| Control | F | 32 | White | None | None | None | None | None |
| Control | M | 20 | White | None | None | None | None | None |
| Control | M | 42 | White | None | None | None | multivit | multivitamin |

| **Dx** | **Sex** | **Age** | **Race** | **Substance use** | **Comorbid Psych Dx** | **Comorbid Med Dx** | **Medications (as reported)** | **Medications (generic name)** |
| --- | --- | --- | --- | --- | --- | --- | --- | --- |
| SZ | M | 32 | White | None | None | Tardive Dyskinesia, past: Hernia | Li 3000, Haldol 3, Ativan 1, vit E | lithium 3000 [this is more likely to be lithium carbonate 300 mg], haloperidol 3, lorazepam 1, vitamin E |
| SZ | F | 50 | White | Lifetime Cannabis abuse | None | None | Trilafon 8 or 16, Artane 4, Navane 30, Zoloft 200, trazodone 100, Ativan prn | perphenazine 8 or 16, trihexyphenidyl 4, thiothixene 30, sertraline 200, trazodone 100, lorazepam prn |
| SZ | F | 57 | White | None | None | Hypercholesterolemia | Risperdal 6, Effexor 150, Tegretol 400, Ativan 1.5 | risperidone 6, venlafaxine 150, carbamazepine 400, lorazepam 1.5 |
| SZ | F | 46 | White | Lifetime Alcohol, Cannabis abuse | Current Panic disorder | Hypothyroid | Klonopin .5, Trileptal 300, Levoxyl 112 mcg, Colace 100, Celexa 60, Clozaril 125, Ditropan 5 | clonazepam .5, oxcarbazepine 300, levothyroxine 112 mcg, docusate 100, citalopram 60, clozapine 125, oxybutynin 5 |
| SZ | M | 43 | White | Lifetime Cannabis dependence | Current AWOPD | None | Clozaril 400, Depakote 1250, Cogentin 2, Lipitor 10 | clozapine 400, valproate 1250, benztropine 2, atorvastatin 10 |
| SZ | M | 50 | White | Lifetime Alcohol, Cannabis,  Hallucinogen -PCP abuse | None | Hypertension | Clozaril 500mg, Klonopin 2mg, Navane 20mg, Paxil 10mg, Neurontin 20mg, BP med | clozapine 500mg, clonazepam 2mg, thiothixene 20mg, paroxetine 10mg, gabapentin 20mg, BP med |
| SZ | M | 31 | White | Current Cannabis dependence | Lifetime Depressive disorder NOS | Raynaud's | Lamictal 150mg, Risperdal 2mg | lamotrigine 150mg, risperidone 2mg |
| SZ | F | 36 | White | None | None | Sleep Apnea, Diabetes, Narcolepsy | Risperdal, Wellbutrin, Cogentin, Ativan | risperidone, bupropion, benztropine, lorazepam |
| SZ, disorg | M | 48 | White | Lifetime Cannabis dependence | None | past: Cryptorchism | Li carbonate 1800, Abilify 15, Klonopin 1 | lithium carbonate 1800, aripiprazole 15, clonazepam 1 |
| SZ, paranoid | M | 27 | not given | Lifetime Alcohol abuse | Current Anxiety NOS | Wolf-Parkinson White Syndrome, Lactose Intolerant | Abilify 30, Clozaril 225, Effexor 225, Haldol 5prn, Metoprolol 100, Ativan 2prn, Prilosec 20, Robinul 2, Maalox 30ml, milk of magnesia | aripiprazole 30, clozapine 225, venlafaxine 225, haloperidol 5prn, metoprolol 100, lorazepam 2prn, omeprazole 20, glycopyrrolate 2, aluminum hydroxide/magnesium hydroxide 30ml, magnesium hydroxide |
| SZ,  undiff | M | 32 | White | Lifetime Alcohol abuse, Cannabis dependence | None | Eczema, Asthma | albuterol | albuterol |
| SZ | M | 44 | White | Lifetime Alcohol, Cannabis, Cocaine dependence | None | None | Clozaril 700, Invega, atenolol, simvastatin | clozapine 700, paliperidone, atenolol, simvastatin |
| SZ | M | 40 | White | Lifetime Alcohol, Cannabis abuse; Current Alcohol  dependence, Cannabis abuse | None | Sleep Apnea | Klonopin 4mg, Abilify 5mg, trazodone 100mg | clonazepam 4mg, aripiprazole 5mg, trazodone 100mg |
| SZ, undiff | M | 21 | White | Lifetime Cannabis abuse | None | past: Cellulitis | clozapine 325, multivit | clozapine 325, multivitamin |
| SZ | F | 67 | White | None | None | Arthritis | Ativan, Celexa 40mg, Trilafon 4mg, Neurontin | lorazepam, citalopram 40mg, perphenazine 4mg, gabapentin |
| SZ | M | 47 | White | Lifetime Alcohol, Cannabis dependence; Current Cannabis dependence | Current Panic disorder | None | Risperdal 8mg, Seroquel 50mg, fluoxetine 40mg | risperidone 8mg, quetiapine 50mg, fluoxetine 40mg |
| SZ | M | 52 | Black | Lifetime Alcohol dependence | None | past: Prostate Cancer; Collapsed Lung | None | None |
| SZ | M | 22 | White | Current Cannabis dependence; Unknown Alcohol, Hallucinogen-PCP | None | None | Li 300mg x 4, Klonopin 6mg tid, Abilify 30mg, Strattera 125mg, Buspar | lithium 300mg x 4, clonazepam 6mg tid, aripiprazole 30mg, atomoxetine 125mg, buspirone |
| SZ | M | 43 | White | Lifetime Alcohol dependence; Cannabis abuse | None | past: Hypertension, Hyperlipidemia, Possible h/o Seizure | Zyprexa 10mg, Atenolol 20mg, simvastatin 20mg | olanzapine 10mg, atenolol 20mg, simvastatin 20mg |
| SZ | F | 68 | White | Lifetime Cannabis, Sedative, Stimulant dependence | Current Specific phobia | past: Meningeal brain tumor on left temporal region | levothyroxine 125ug | levothyroxine 125mcg |
| SZ | M | 53 | White | None | None | Pressure headaches, Arthritis | Zyprexa 10mg, Zyprexa 2.5mg prn | olanzapine 10mg, olanzapine 2.5mg prn |
| SZ | M | 42 | White | Lifetime Stimulant abuse | None | Schwannoma, past: Chronic back pain, Bundle branch block; NSAID-Induced Ulcers | clozapine 825mg, Valium 20mg, Inderal 10mg, Zocor 20mg, Prilosec, Detrol LA 4mg, Lidoderm 5% patch, Robinul 2mg, vitamin E, vit D, vit C, multivit, Percocet 5/325 q.6h prn | clozapine 825mg, diazepam 20mg, propranolol 10mg, simvastatin 20mg, omeprazole, tolterodine 4mg, lidocaine 5% patch, glycopyrrolate 2mg, vitamin E, vitamin D, vitamin C, multivitamin, oxycodone/acetaminophen 5/325 q.6h prn |
| SZ | M | 50 | White | Lifetime Alcohol, Cannabis dependence; Current Alcohol dependence | None | Recent fall: Broken Clavicle, COPD, Arthritis, Dizziness | Thorazine 100, Prolixin 15, Inderal 60, Ativan prn 1, Haldol prn 5, pravastatin 40, Spiriva 18mcg, Prilosec 20, nicotine 14, ibuprofen 600 | chlorpromazine 100, fluphenazine 15, propranolol 60, lorazepam prn 1, haloperidol prn 5, pravastatin 40, tiotropium 18mcg, omeprazole 20, nicotine 14, ibuprofen 600 |
| SZ | M | 39 | White | None | Lifetime Panic disorder | None | Geodon 40, trazodone 300, Klonopin 2, Androgel 1g, Propecia 1mg, Norvasc 10 | ziprasidone 40, trazodone 300, clonazepam 2, testosterone gel 1g, finasteride 1mg, amlodipine 10 |
| SZ | F | 39 | White | Lifetime Alcohol, Polydrug dependence | Lifetime Bulimia | None | levothyroxine 50, Topamax 100, Haldol 30, Benadryl 50, Rozerem 8, Tegretol 600 | levothyroxine 50, topiramate 100, haloperidol 30, diphenhydramine 50, ramelteon 8, carbamazepine 600 |
| SZ | M | 42 | Black | None | None | None | Zyprexa 30mg | olanzapine 30mg |
| SZ, paranoid | M | 28 | White | None | None | None | Li 900mg, perphenazine | lithium 900mg, perphenazine |
| SZ | M | 23 | White | Current Cannabis dependence | None | Hypoglycemia, past: Gluteal Cystic Hygroma, ER for Foot Infection | Abilify 10mg, Ativan 1prn | aripiprazole 10mg, lorazepam 1prn |
| SZ,  undiff | M | 53 | White | Lifetime Alcohol abuse | None | Overweight | Zyprexa 30, Zyprexa 5prn, Depakote 2250, Paxil 60, Ativan 3mg prn | olanzapine 30, olanzapine 5prn, valproate 2250, paroxetine 60, lorazepam 3mg prn |
| SZ | M | 23 | White | None | None | None | Risperdal 120 | risperidone 120 |
| SZ | M | 51 | White | None | None | Anemia, GERD,  past: Pilonidal Cyst | clozapine 350, Zyprexa 10, Hexum 40, aspirin 81, Ativan 1mg prn | clozapine 350, olanzapine 10, hexum 40, aspirin 81, lorazepam 1mg prn |
| SZ | M | 23 | Asian/  White | None | None | None | selegiline 12, clonazepam 2, zolpidem 12.5, l-methylfolate 15, Aricept 23, Cytomel (T3) 50 | selegiline 12 mg, clonazepam 2 mg, zolpidem 12.5 mg, l-methylfolate 15 mg, donepezil 23 mg, liothyronine 50 mcg |
| SZ | M | 19 | White | Lifetime Alcohol, Cannabis, Hallucinogen dependence,  Sedative, Hypnotic, Anxiolytic abuse; Current Cannabis dependence | None | Chronic Migraines, Stomach Problems | Cogentin 0.5, Seroquel 400, Ambien 20, Seroquel 400 prn, Inderal 120, multivit | benztropine 0.5, quetiapine 400, zolpidem 20, quetiapine 400 prn, propranolol 120, multivitamin |
| SZA | M | 58 | Black | None | None | None | Zyprexa, Depakote | olanzapine, valproate |
| SZ | M | 24 | Asian | Lifetime: Alcohol abuse; cannabis dependence | None | None | Abilify 10, Ativan 2, propranolol prn bid 20 | aripiprazole 10, lorazepam 2, propranolol prn bid 20 |
| SZ | M | 19 | White | None | None | Chest pains | risperidone 2 prn, clonazepam prn, metformin 500 | risperidone 2 prn, clonazepam prn, metformin 500 |
| Psychosis NOS | M | 22 | White | Not given | Not given | Not given | Not given | Not given |
| SZ | M | 23 | White | Lifetime Cannabis abuse | None | None | Li 1500mg | lithium 1500mg |
| SZ | M | 18 | White | None | Current Social phobia, OCD | None | Li 900mg qhs, clozapine 200mg qhs, Ativan 0.5mg bid | lithium 900mg qhs, clozapine 200mg qhs, lorazepam 0.5mg bid |
| SZform/  SZA | M | 19 | Black | Lifetime Cannabis abuse | None | None | Celexa 20mg, risperidone 3mg at bedtime | citalopram 20mg, risperidone 3mg at bedtime |
| SZA | M | 44 | White | None | None | Diabetes | Li carbonate 300mg + 600mg, clozapine 125mg bid, Depakote 500 & 700mg, simvastatin, multivit | lithium carbonate 300mg + 600mg, clozapine 125mg bid, valproate 500 & 700mg, simvastatin, multivitamin |
| SZA | F | 29 | White | None | Lifetime Panic disorder | None | Prozac 20mg, Abilify 20mg, Lamictal 250mg, Provigil 100mg, St. John's wort, multivit | fluoxetine 20mg, aripiprazole 20mg, lamotrigine 250mg, modafinil 100mg, St. John's wort, multivitamin |
| SZA, bipolar | M | 33 | White | Lifetime Cannabis abuse | None | Asthma | Li 1800, aripiprazole 40, oxcarbazepine 1500, citalopram 20, lorazepam 1 prn | lithium 1800, aripiprazole 40, oxcarbazepine 1500, citalopram 20, lorazepam 1 prn |
| SZA | F | 20 | White | Lifetime Cannabis abuse | None | past: Chronic Migraines | Celexa 20, Risperdal 3, Ativan 1 prn | citalopram 20, risperidone 3, lorazepam 1 prn |
| SZA | M | 45 | White | None | Lifetime Panic disorder w/o Agoraphobia | None | Effexor 150, ability 15, Ativan 1 | venlafaxine 150, aripiprazole 15, lorazepam 1 |
| SZA | F | 25 | White | None | Lifetime Social phobia | Macroprolactinoma. Galactorrhea. Hypertension | risperidone 4, Lamictal 150, Paxil 30, trazodone prn 50, aripiprazole | risperidone 4, lamotrigine 150, paroxetine 30, trazodone prn 50, aripiprazole |
| SZA | M | 22 | White | Lifetime Alcohol, Cocaine, Hallucinogen abuse; Cannabis dependence | None | None | Li 600mg, aripiprazole 30mg, olanzapine 25mg, lorazepam 0.5mg, benztropine 25mg | lithium 600mg, aripiprazole 30mg, olanzapine 25mg, lorazepam 0.5mg, benztropine 25mg |
| SZA | M | 41 | White | None | None | Asthma | Abilify 20mg, Venlafaxine XR 150mg, Zafirlukast 40mg, aspirin 81mg | aripiprazole 20mg, venlafaxine XR 150mg, zafirlukast 40mg, aspirin 81mg |
| SZA | F | 38 | White | None | Current Depressive disorder NOS, Lifetime Social phobia | Hypothyroidism, Breast Reduction | clozapine 300mg, citalopram 40mg, levothyroxine 50mcg, lorazepam 1mg, glycopyrrolate 1mg, vit D 800 | clozapine 300mg, citalopram 40mg, levothyroxine 50mcg, lorazepam 1mg, glycopyrrolate 1mg, vitamin D 800 |
| SZA | M | 27 | White | None | None | Lyme Disease | risperidone 3mg, clonazepam 0.5mg | risperidone 3mg, clonazepam 0.5mg |

| **Dx** | **Sex** | **Age** | **Race** | **Substance Use** | **Comorbid Psych Dx** | **Comorbid Med Dx** | **Medications (as reported)** | **Medications (generic name)** |
| --- | --- | --- | --- | --- | --- | --- | --- | --- |
| BPI | M | 47 | White | None | Current: OCD, PTSD, GAD, Agoraphobia, w/o PD | None noted | Advair 100/50 BID, Ventolin (inhaler) 2 puffs | fluticasone/salmeterol 100/50 BID, albuterol (inhaler) 2 puffs |
| BPII | M | 54 | White | Lifetime Alcohol abuse, Unknown Cannabis | None | None | Li 1500 mg, Lamictal 300mg, Provigil 200mg, Topamax 100mg | lithium 1500 mg, lamotrigine 300mg, modafinil 200mg, topiramate 100mg |
| BPI | F | 35 | White | None | None | Genital Herpes | Risperdal 0.5, Lamictal .75, Yasmin BCP | risperidone 0.5, lamotrigine .75, drospirenone/ethinyl estradiol BCP |
| BPI | M | 19 | White | Lifetime Cannabis abuse, Current Cannabis dependence | None | None | risperidone, Ativan | risperidone, lorazepam |
| BPI | F | 43 | White | None | Lifetime Panic disorder | None noted | naproxen bid, Singulair 10mg hs, Advair 50/50 bid, Flovent 225/50 bid, albuterol prn, Protonix 10 am, Cymbalta 60am, Seroquel 75-100 hs | naproxen bid, montelukast 10mg hs, fluticasone/salmeterol 50/50 bid, fluticasone 225/50 bid, albuterol prn, pantoprazole 10 am, duloxetine 60am, quetiapine 75-100 hs |
| BPI | M | 25 | White | Lifetime: Alcohol dependence, Cannabis abuse | None | Controlled Hypothyroid, Giardia Parasite (treated), Ulcerative colitis | Ativan 3, Risperdal 6, Lamictal 200, mercaptopurine 125, Asacol 3200, Haldol 5 prn | lorazepam 3, risperidone 6, lamotrigine 200, mercaptopurine 125, mesalamine 3200, haloperidol 5 prn |
| BPI | M | 58 | White | None | Current PTSD | Sleep Apnea; Hypertension | Li 1500, omeprazole 20, Lipitor 40, Lisinopril 10, Actos 30, Lasix 40, Neurontin 600, Coumadin 12, Geodon 1200, Klonopin 2 | lithium 1500, omeprazole 20, atorvastatin 40, lisinopril 10, pioglitazone 30, furosemide 40, gabapentin 600, warfarin 12, ziprasidone 1200, clonazepam 2 |
| BPII | M | 43 | White | Lifetime Alcohol, Polydrug dependence | None | None | Lamictal 150mg, Lexapro 10mg, Provigil 200mg, Risperdal 5mg, Depakote 2250mg | lamotrigine 150mg, escitalopram 10mg, modafinil 200mg, risperidone 5mg, valproate 2250mg |
| BPI | M | 20 | White | Lifetime Alcohol dependence, Cannabis abuse; Current Alcohol, Cannabis dependence | Current OCD | None | Li 1200mg, Risperdal 4mg | lithium 1200mg, risperidone 4mg |
| BPI | F | 35 | White | None | Current Specific phobia | None | Neurontin 300mg tid, Zyprexa 10mg, Ativan 1mg, Seasonale, Prozac, vits, fish oil, naltrexone | gabapentin 300mg tid, olanzapine 10mg, lorazepam 1mg, levonorgestrel/ethinyl estradiol, fluoxetine, vitamins, fish oil, naltrexone |
| BPI | F | 27 | White | None | None | None | Li 450mg, Wellbutrin | lithium 450mg, bupropion |
| BPI | F | 40 | White | Lifetime Alcohol, Cannabis dependence | None | Cardiomyopathy | Seroquel 25mg, Klonopin 1mg, Prilosec 2mg | quetiapine 25mg, clonazepam 1mg, omeprazole 2mg |
| BPI | F | 39 | White | None | Lifetime OCD, Current GAD | Hypertension, Low Potassium | Seroquel 700, Trileptal 1500, Ativan 1, HCTZ 25 | quetiapine 700, oxcarbazepine 1500, lorazepam 1, hydrochlorothiazide 25 |
| BPI | F | 41 | Asian | None | None | None | Depakote, naltrexone | valproate, naltrexone |
| BPI | M | 55 | White | None | None | past: 2 Pulmonary Eboli | Seroquel 200mg, vitamins, ECT | quetiapine 200mg, vitamins, ECT |
| BPI | M | 36 | White | Lifetime Cannabis dependence | Current GAD | None | Trileptal 600 bid, Cymbalta 60, Klonopin 2 | oxcarbazepine 600 bid, duloxetine 60, clonazepam 2 |
| BPI | M | 46 | White | None | None | past: Migraines, Hypertension | Ativan 2mg, Zyprexa 15mg, Depakote 2000mg | lorazepam 2mg, olanzapine 15mg, valproate 2000mg |
| BPI | M | 62 | White | Lifetime Alcohol dependence | Current Social phobia, Specific phobia | Hypertension, Arthritis, GERD | Cymbalta 90, Seroquel 800, Topamax 75, atenolol 25, Feldene 1040, Naprosyn 1000, Flonase, Prilosec | duloxetine 90, quetiapine 800, topiramate 75, atenolol 25, piroxicam 1040, naproxen 1000, fluticasone nasal, omeprazole |
| BPI | M | 28 | White | Lifetime Alcohol, Cannabis, Hallucinogen -PCP dependence | None | None | risperidone 2, Lamictal 300 | risperidone 2, lamotrigine 300 |
| BPI | M | 25 | White | None | None | None | Li 1800, Zyprexa 20, Zyprexa prn 10, Ativan 1-3, propranolol 400, multivit | lithium 1800, olanzapine 20, olanzapine prn 10, lorazepam 1-3, propranolol 400, multivitamin |
| BPI | M | 43 | White | None | None | None | Clozaril 175, Depakote 750, Lipitor 40, Propecia 1mg, Advair, O3FA | clozapine 175, valproate 750, atorvastatin 40, finasteride 1mg, fluticasone/salmeterol, omega-3 fatty acids |
| BPI w/ psychosis | M | 41 | White | Lifetime Alcohol, Cannabis, Stimulant, Hallucinogen abuse | None | Otitis Media, past: Heart Murmur, Kidney Stones | Li 900, risperidone 2, Seroquel 50 prn, Ativan 1prn | lithium 900, risperidone 2, quetiapine 50 prn, lorazepam 1prn |
| BPI | M | 23 | White | Lifetime Alcohol, Cannabis dependence; Current Cannabis dependence | None | past: decreased Renal Function | Li ER 600+1200mg, risperidone 1+3mg (plus up to 2x/day 1mg prn), Lamictal 100+100mg, Zyprexa 20mg | lithium ER 600+1200mg, risperidone 1+3mg (plus up to 2x/day 1mg prn), lamotrigine 100+100mg, olanzapine 20mg |
| BPI | M | 32 | White | Lifetime Alcohol, Cannabis dependence | None | past: Complex Partial Seizures, Chlamydia; Gonorrhea | Li ER (900+600), Geodon | lithium ER (900+600), ziprasidone |
| BPI | M | 40 | White | None | Current Social phobia | None | Li 900mg | lithium 900mg |
| BPI/SZA* | F | 39 | White | Lifetime Cocaine dependence | Social phobia | None | no meds | no meds |
| BPI | M | 31 | White | Lifetime Cannabis dependence | Lifetime Panic disorder | Chronic Migraines | Abilify 5, Klonopin 1 | aripiprazole 5, clonazepam 1 |
| BPI | M | 27 | White | Lifetime Alcohol dependence | None | Staph infection | Risperdal 2mg, biotin, Zyrtec, Claritin, sulfameth | risperidone 2mg, biotin, cetirizine, loratadine, sulfamethoxazole |
| BPI |  | 26 | other | Lifetime Alcohol abuse, Cannabis dependence; Current Cannabis dependence | None | past: Toxic Epidermal Necrolysis, Inguinal hernia, Asthma, Skin graft on foot | Li 600 + 900=1500, Zyprexa 20, multivit | lithium 600 + 900=1500, olanzapine 20, multivitamin |
| BPI | M | 55 | White | None | None | past: Hypertension | aspirin 325, propranolol 20, Depakote 2000, simvastatin 20, Seroquel 150 | aspirin 325, propranolol 20, valproate 2000, simvastatin 20, quetiapine 150 |
| BPI | F | 43 | White | None | None | None | olanzapine 15mg, lorazepam 2mg, Depakote 1000mg | olanzapine 15mg, lorazepam 2mg, valproate 1000mg |
| BPI | M | 25 | White | Lifetime Alcohol, Cannabis, dependence; Sedative, Hypnotic, Anxiolytic abuse | None | None | None | None |
| BPI | F | 22 | White | Lifetime Cannabis dependence | Not reported | Shingles, Chronic Migraines | Li 1200, Lamictal 150, Seroquel 200 | lithium 1200, lamotrigine 150, quetiapine 200 |
| BPI | M | 20 | White | Lifetime Cannabis abuse | Current Panic disorder | Weight loss, Chronic Migraines | Li 900, Zyprexa 20, Ativan 1 prn, multivit | lithium 900, olanzapine 20, lorazepam 1 prn, multivitamin |
| BPI/SZA* | M | 30 | mixed | None | None | None | Abilify 20, Klonopin 2, Trileptal 1200, Lopressor 50 | aripiprazole 20, clonazepam 2, oxcarbazepine 1200, metoprolol 50 |
| BPI NOS | M | 58 | White | None | Panic DO | Hypercholesterolemia | loratadine 400mg, Klonopin 0.15 qits 0.5 qam, Ambien 10mg, atorvastatin 20mg, aspirin 81mg, O3FA 1000 mg, Tramadol 50mg prn | loratadine 400mg, clonazepam 0.15 qits 0.5 qam, zolpidem 10mg, atorvastatin 20mg, aspirin 81mg, omega-3 fatty acids 1000 mg, tramadol 50mg prn |
| BPI NOS | M | 59 | Black | None | PTSD, AWOPD | Type II Diabetes, Hyperlipidemia, Hypertension | labetalol 600mg bid, metformin 1000mg bid, simvastatin 20mg qd, aspirin 81mg qd, glyburide 5mg bid, Cozaar 100mg qd, HCTZ 25mg qd | labetalol 600mg bid, metformin 1000mg bid, simvastatin 20mg qd, aspirin 81mg qd, glyburide 5mg bid, losartan 100mg qd, hydrochlorothiazide 25mg qd |
| BPI | F | 33 | White | None | Current AWOPD, Social phobia, OCD, GAD | Fatigue, Diarrhea | Seroquel 25mg prn, lorazepam .5mg prn, Lamictal 150mg, Diflunisal 1000mg, venlafaxine ER, hyoscyamine sulfate 0.125mg prn, sumatriptan 50mg prn, Trinessa BCP | quetiapine 25mg prn, lorazepam .5mg prn, lamotrigine 150mg, diflunisal 1000mg, venlafaxine ER, hyoscyamine sulfate 0.125mg prn, sumatriptan 50mg prn, norgestimate/ethinyl estradiol BCP |
| BPI | F | 69 | White | None | None | Macular Degeneration | Ritalin 60mg qam, triamterene/HCTZ 75/50mg qam, Effexor XR 300mg qam, Trileptal 600mg bid, Seroquel 50mg | methylphenidate 60mg qam, triamterene/hydrochlorothiazide 75/50mg qam, venlafaxine XR 300mg qam, oxcarbazepine 600mg bid, quetiapine 50mg |
| BPI | M | 24 | White | Lifetime Alcohol, Cannabis abuse | None | None | None | None |
| BPI | M | 22 | Black | Lifetime Cannabis abuse | None | None | Li 1200 | lithium 1200 |
| BPI | F | 20 | White | Lifetime Cannabis, Alcohol abuse | None | None | Li 600, bupropion 300, lorazepam 0.5, Beyaz BCP | lithium 600, bupropion 300, lorazepam 0.5, drospirenone/ethinyl estradiol/levomefolate BCP |
| BPI | M | 21 | White | Not reported | Not reported | None | Not reported | Not reported |
| BPI | F | 37 | White | Lifetime Alcohol, Cannabis, Stimulant abuse | Current OCD, Personality disorder | Kidney Infection, Seizure Disorder | Lamictal 500, Lexapro 5, Klonopin 1, fish oil 900 | lamotrigine 500, escitalopram 5, clonazepam 1, fish oil 900 |
| BPI | M | 41 | White | Lifetime Alcohol abuse; Cannabis dependence | Lifetime MDD | None | Li 1050mg | lithium 1050mg |
| BPI | M | 22 | Asian | None | Current OCD | None | Li carbonate ER 1200mg, risperidone 1mg, fish oil | lithium carbonate ER 1200mg, risperidone 1mg, fish oil |
| BPI | M | 24 | White | Lifetime Alcohol dependence, Cannabis abuse | None | None | Li 900, risperidone 4mg, lorazepam 2mg | lithium 900, risperidone 4mg, lorazepam 2mg |
| BPI | M | 24 | White | None | None | None | Depakote ER 1250mg | valproate ER 1250mg |
| BPI | M | 21 | White | Lifetime Alcohol dependence, Cannabis abuse | None | None | olanzapine 10mg bid, gabapentin 300mg 4x/day, Depakote 500x4 at night | olanzapine 10mg bid, gabapentin 300mg 4x/day, valproate 500x4 at night |
| BPI | M | 27 | White | Lifetime Cannabis dependence | None | None | Li 900 | lithium 900 |

*Later SCID gave Dx change to SZA

| **Dx** | **Sex** | **Age** | **Race** | **Substance Use** | **Comorbid Psych Dx** | **Comorbid Med Dx** | **Medications (as reported)** | **Medications (generic name)** |
| --- | --- | --- | --- | --- | --- | --- | --- | --- |
| MDD | M | 51 | White | None | Current Anxiety NOS | Hypertension, Hypercholesterolemia, GERD, Sleep Apnea | Seroquel 100 mg, Lorazepam 1 mg, Remeron 15 mg, Lisinopril 40 mg, Simvastatin 20 mg, Omeprazole 40 mg, Ibuprofen 400 mg, multivit | quetiapine 100 mg, lorazepam 1 mg, mirtazapine 15 mg, lisinopril 40 mg, simvastatin 20 mg, omeprazole 40 mg, ibuprofen 400 mg, multivit |
| MDD | F | 41 | White | None | None | Sleep Apnea | Abilify 7.5mg, Thorazine 25mg, Colace 100mg, Seroquel 50mg, Cytomel 15ug, Vistaril 50mg, Klonopin 1mg, Maalox, Tylenol | aripiprazole 7.5mg, chlorpromazine 25mg, docusate 100mg, quetiapine 50mg, liothyronine 15mcg, liothyronine 50mg, clonazepam 1mg, magnesium hydroxide, acetaminophen |
| MDD | F | 52 | White | None | None | Renal Cell Carcinoma, Abnormal Thyroid Function | Li carbonate 300mg, Wellbutrin, trazodone 100mg prn, Neurontin 600 qhs, Klonopin | Li carbonate 300mg, bupropion, trazodone 100mg prn, gabapentin 600 qhs, clonazepam |
| MDD | F | 19 | Black | None | Current Social phobia | Migraines | Celexa 30mg, trazodone 25mg, Ativan, Colace, Benadryl, Maalox, Tylenol | citalopram 30mg, trazodone 25mg, lorazepam, docusate, diphenhydramine, magnesium hydroxide, acetaminophen |
| MDD | M | 19 | White | None | Current Anxiety NOS | None | Wellbutrin 75mg bid, olanzapine 5mg, Ativan, Maalox, Tylenol, Dulcolax | bupropion 75mg bid, olanzapine 5mg, lorazepam, aluminum hydroxide/magnesium hydroxide, acetaminophen, bisacodyl |
| MDD | M | 32 | White | Lifetime Opioid dependence | None | Diabetes, Gastroparesis, Hypothyroidism, Neuropathy | Lyrica 300, Effexor 300, Zofran 8x3, Lamictal 200, Abilify 5, Wellbutrin 100, methadone 30, Synthroid 100mcg, NicoDerm, insulin, Ativan prn, Benadryl prn, multivit | pregabalin 300, venlafaxine 300, ondansetron 8x3, lamotrigine 200, aripiprazole 5, bupropion 100, methadone 30, levothyroxine 100mcg, nicotine patch, insulin, lorazepam prn, diphenhydramine prn, multivit |
| MDD | F | 22 | White | None | None | None | Zoloft 100mg qam, Ativan .25 prn | sertraline 100mg qam, lorazepam .25 prn |
| MDD  w/ psy | M | 25 | SE Asian | None | Current Anxiety NOS | Chronic "Tension Headaches" | Zyprexa 15, Celexa 20, Zyprexa 2.5 prn, baclofen 10 prn | olanzapine 15, citalopram 20, olanzapine 2.5 prn, baclofen 10 prn |
| MDD | F | 36 | White | None | Current GAD | Migraines, past: Bruxism | Cymbalta, Trazodone, Klonopin | duloxetine, trazodone, clonazepam |
| MDD | F | 33 | White | None | Current OCD, GAD | Celiac Disease, Ulcerative Colitis | Li Carbonate 300mg, risperidone 2mg qh, 600mg, lorazepam 0.5mg prn q4h, Sertraline HCl 100mg qd, ethinyl estradiol/norgestimate 1 tab hs, multivit | lithium carbonate 300mg, risperidone 2mg qh, 600mg, lorazepam 0.5mg prn q4h, sertraline 100mg qd, ethinyl estradiol/norgestimate 1 tab hs, multivit |
| MDD | M | 22 | White | None | None | Asthma | albuterol | albuterol |
| MDD | M | 55 | White | None | Current PTSD, GAD | past: In Cardiac Unit | Cymbalta 60, Seroquel, Testim gel, Lamictal, Valium prn, Tenormin, Crestor, aspirin | duloxetine 60, quetiapine, testosterone gel, lamotrigine, diazepam prn, atenolol, rosuvastatin, aspirin |
| MDD | M | 32 | Asian | None | Current Social phobia | Gastroesophageal Reflux | Zoloft 100mg, Vistaril 50mg, Prilosec 20mg | sertraline 100mg, hydroxyzine 50mg, omeprazole 20mg |
| MDD | F | 40 | White | Lifetime Alcohol dependence | Current PTSD | Chronic Migraines, Allergy Induced Asthma, Sleep Apnea | Effexor 225 mg, Risperdal 2 mg, fluticasone spray | venlafaxine 225 mg, risperidone 2 mg, fluticasone spray |
| MDD | F | 21 | White | None | None | Asthma, Hypercholesterolemia, Orthostatic Hypotension (past): TMJ surgery | Wellbutrin 100mg, multivits | bupropion 100mg, multivits |
| MDD | M | 18 | Black | None | Current PTSD | None | Zoloft 25, Ativan 1 mg prn, Zyprexa 2.5 mg prn, Tylenol 650 | sertraline 25, lorazepam 1 mg prn, olanzapine 2.5 mg prn, acetaminophen 650 |
| MDD | M | 30 | White | Lifetime Alcohol, Sedative, Hypnotic, Anxiolytic abuse; Stimulant dependence | Lifetime Social phobia, OCD | None | bupropion 300mg, clomipramine 100mg, clonazepam 3mg | bupropion 300mg, clomipramine 100mg, clonazepam 3mg |
| MDD | F | 26 | White | Current Cannabis dependence | Current Anxiety NOS | None | mirtzapine 30mg | mirtzapine 30mg |
| MDD | F | 27 | White | Lifetime Alcohol abuse | Lifetime PTSD | None | Trilafon 20gm/day, Zoloft 100mg/day | perphenazine 20gm/day, sertraline 100mg/day |
| MDD | M | 24 | White | Lifetime Cannabis dependence | None | None | Celexa 40mg, Abilify 5mg | citalopram 40mg, aripiprazole 5mg |
